# Heme oxygenase-1 expressing omental macrophages as a therapeutic target in ovarian high grade serous carcinoma

**DOI:** 10.1101/2023.07.18.549474

**Authors:** Sarah Spear, Olivia Le Saux, Hasan B. Mirza, Katie Tyson, Jasmine Bickel, Fabio Grundland Freile, Alexandros P. Siskos, Cristina Balcells, Josephine B. Walton, Chloé Woodman, Darren P. Ennis, Nayana Iyer, Carmen Aguirre Hernandez, Yuewei Xu, Pavlina Spiliopoulou, James D. Brenton, Ana P. Costa-Pereira, Hector C. Keun, Evangelos Triantafyllou, James N. Arnold, Iain A. McNeish

## Abstract

Ovarian high grade serous carcinoma (HGSC) remains a disease of poor prognosis that is unresponsive to current immune checkpoint inhibitors. Although PI3K pathway alterations are common in HGSC, attempts to target this pathway have been unsuccessful. We hypothesised aberrant PI3K pathway activation may alter the HGSC immune microenvironment and present a novel targeting strategy. We used both murine models and HGSC patient samples to study the impact of loss of *Pten*, a negative regulator of PI3K pathway signalling. We identified populations of resident macrophages specifically in *Pten* null omental tumours. These macrophages derive from peritoneal fluid macrophages and have a unique gene expression programme, marked by high levels of *HMOX1* expression, the gene for the enzyme heme oxygenase-1. Targeting resident peritoneal macrophages prevents appearance of HMOX1^hi^ macrophages and in doing so reduces tumour growth. Furthermore, direct inhibition of HMOX1 extends survival *in vivo*. HMOX1^hi^ macrophages with corresponding gene expression programmes are also identified in human HGSC tumours and their presence correlates with activated tumoural PI3K pathway/mTOR signalling and poor overall survival in HGSC patients. In contrast, tumours with low number of HMOX1^hi^ macrophages are marked by increased adaptive immune response gene expression. Our data suggest that HMOX1^hi^ macrophages represent a potential therapeutic target and biomarker for poor prognosis HGSC.

## Introduction

High grade serous carcinoma (HGSC), the commonest type of ovarian cancer, remains a disease of poor prognosis, especially for patients whose tumours are classified as having proficient homologous recombination^1^. The immune microenvironment has a strong prognostic effect in HGSC^2^. The presence of intra-epithelial CD8^+^ T cells^3^ and immunoreactive gene expression signatures^4, 5^ both correlate with improved overall survival whilst intra-tumoural Treg are associated with poor survival^6^. However, responses to immune checkpoint inhibitors are poor^7, 8^ with little correlation between response and either platinum-free interval or tumour cell PD-L1 expression^9^. Putative neoantigens can be identified in HGSC^10^, but average mutational burden is low^11^.

PTEN is a tumour suppressor and negative regulator of the PI3K-signalling pathway. Although deleterious single nucleotide variants and deletions in *PTEN* are rare in HGSC^4^, inactivation through complex re-arrangements is more frequent^12^, and complete (13–59%) or partial (13–55%) PTEN protein loss is common^13–16^, suggesting transcriptional dysregulation. Furthermore, genomic alterations in the PI3K/Ras signalling pathway occur in 45% of HGSC^4^ and enhanced PI3K pathway signalling is observed in the presence of PTEN protein expression^17^. Thus, the PI3K pathway represents an important therapeutic target in HGSC. However, clinical trials of small molecule inhibitors have been largely negative so far^18–20^ and there remains a need to identify effective therapeutic strategies for these tumours.

We hypothesised that loss of PTEN and activated PI3K signalling supports HGSC growth in part through an interaction with the tumour microenvironment. Using murine models, we identified that PTEN loss drives expansion of a resident macrophage population in omental tumours, marked by high expression of the enzyme heme-oxygenase 1 (HMOX1), that likely derives from peritoneal fluid macrophages. We demonstrated that similar HMOX1^hi^ macrophages can be identified in human HGSC samples and are associated with aberrant PI3K pathway activity and poor survival. Finally, we also show that targeting of this population of macrophages may have therapeutic potential in HGSC.

## Results

To address how PTEN loss influences HGSC growth in a controlled system, we utilised matched ID8 cells with inactivating mutations in *Trp53* alone or both *Trp53* and *Pten* that we generated previously^21, 22^. Using multiple separate clones, we confirmed that *Trp53*^-/-^;*Pten*^-/-^ ID8 cells lead to significantly shortened survival compared to *Trp53*^-/-^ following intraperitoneal injection (Fig. 1A). *Pten* deletion, as previously^23^, did not decrease the *in vitro* doubling time in 2D high-attachment conditions across multiple clones (Fig. 1B, C), including those with an additional *Brca2* mutation (Fig. 1D), and under low serum and serum-starvation conditions (Fig. 1E; Fig. S1A), suggesting that enhanced intraperitoneal growth was not tumour-cell intrinsic.

**Figure 1:**
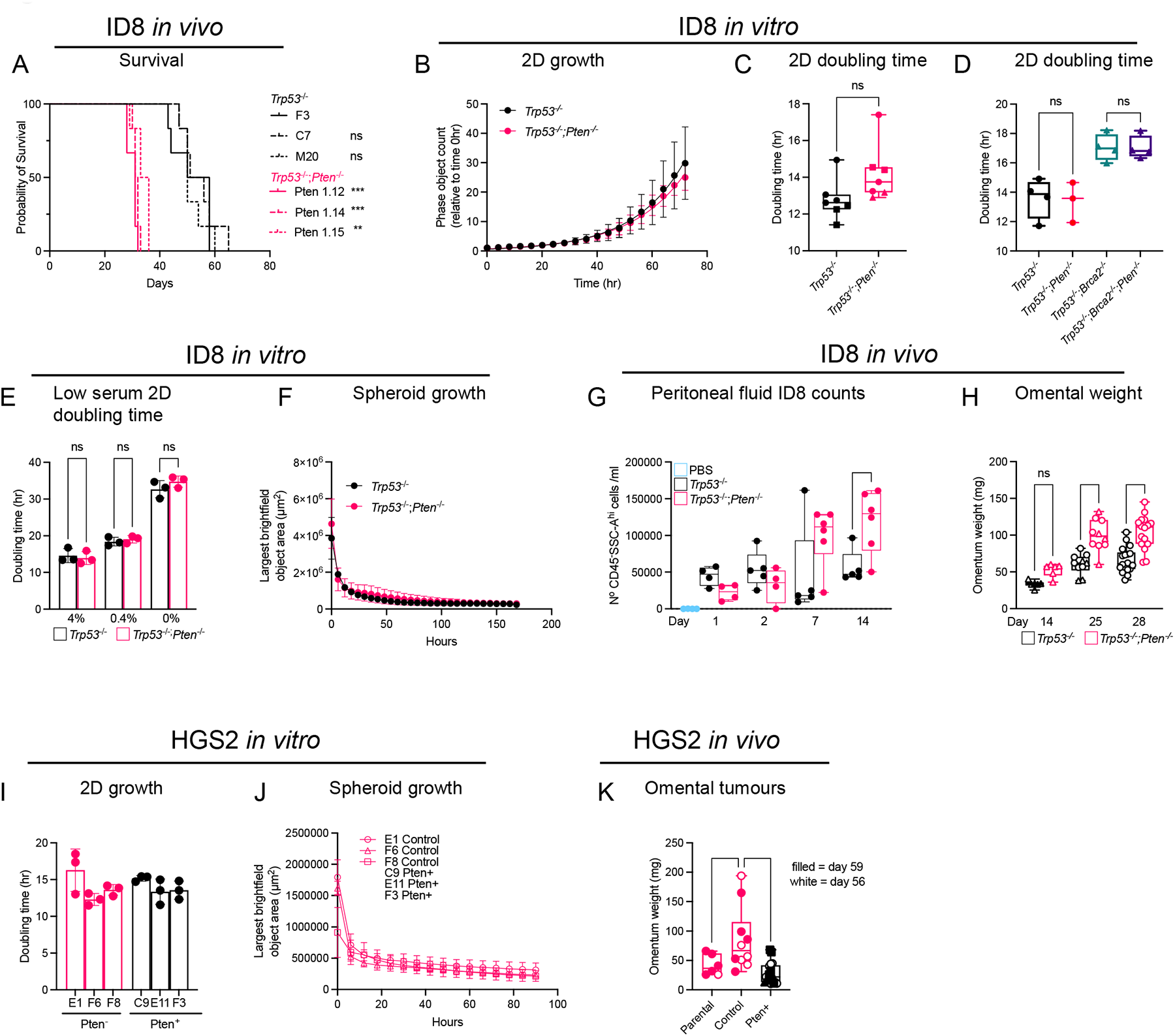
*Pten* null cells are dependent on a tumour microenvironment for accelerated tumour growth. **A)** Survival curve for mice injected with ID8 *Trp53*^-/-^ (clones F3, M20 and C7), and *Trp53*^-/-^;*Pten*^-/-^ (clones Pten1.12, Pten1.14, and Pten1.15), n=6 per clone. Statistical significance was tested using the Log-rank (Mantel-Cox) test. **B)** ID8 *Trp53*^-/-^ (F3) and *Trp53*^-/-^;*Pten*^-/-^ (Pten1.14) cells grown in flat high-attachment plates were imaged every 4hr over 72 hr. Each data point represents the average of 3-4 technical replicates per clone and 4 images per replicate. Data were generated as phase object count per well normalised to first scan (“0 hr”). Representative data from experiment shown. **C)** Mean doubling time of ID8 cells grown in **B)** conditions for 72hr. Each data point represents a clone grown at a different passage or different clone; an average of 3-4 wells, and 4 images per well were used to generate each data point. Clones plated as follows *Trp53*^-/-^ ID8-F3 (circle), ID8-M20 (square), and *Trp53*^-/-^;*Pten*^-/-^, ID8-F3; Pten1.14 (circle), ID8-F3;Pten1.15 (triangle). Significance was tested by an unpaired t-test. **D)** Mean doubling time of ID8 subclones grown in 2D under same conditions as in **C)**, clones used were *Trp53*^-/-^ (F3), *Trp53*^-/-^;*Pten*^-/-^ (Pten1.14), *Trp53*^-/-^;*Brca2*^-/-^ (Brca2 2.14) and *Trp53*^-/-^;*Brca2*^-/-^ *;Pten*^-/-^ (Brca2.14 Pten22). Statistical significance was tested using an ordinary one-way ANOVA, with Šidák’s multiple comparison test on selected pairs. **E)** ID8 *Trp53*^-/-^ (F3), *Trp53*^-/-^;*Pten*^-/-^ (Pten1.14) cells were seeded and grown in 4%, 0.4% or 0% FBS for up to 72hr and the doubling time was calculated as in **C)**. Each symbol represents the average of technical triplicates, performed over 3 passages (P1, P4 and P5). Statistical significance was tested using an ordinary one-way ANOVA, with Šidák’s multiple comparison test on selected pairs. **F)** ID8 clones *Trp53*^-/-^ (F3), *Trp53*^-/-^;*Pten*^-/-^ (Pten1.14) were grown in low-attachment u-bottomed plates for 168 hr. Each symbol represents the average of technical triplicates, repeated on two passages. The largest brightfield object area (µm^2^) per image was quantified and shown over time. **G)** Mice were injected with either PBS, *Trp53*^-/-^ (F3) or *Trp53*^-/-^;*Pten*^-/-^ (Pten1.14) ID8 cells on day 0 and a peritoneal lavage was performed on days 1, 2, 7, and 14. The ID8 cell count was estimated by flow cytometry (gated on as CD45-, SSC-A^hi^, Live). Each point represents an individual mouse. Statistical significance was tested using an ordinary one-way ANOVA, with Šidák’s multiple comparison test on selected pairs. **H)** Mice were injected with *Trp53*^-/-^ (F3) or *Trp53*^-/-^;*Pten*^-/-^ (Pten1.14) ID8 cells on day 0 and the omental tumours harvested at days 14, 25 and 28. Tumour weights shown are pooled from a several experiments, including control groups from other studies. Each point represents an individual mouse. Triangles indicate mice received artificial sweetener in their drinking water for 14 days prior to ID8 injection. Significance was tested by an ordinary one-way ANOVA, with Šidák’s multiple comparison test on selected pairs. **I)** Mean doubling times of HGS2 lentivirus-transduced clones grown for 72 hr in the same conditions as in **C)**. Each circle represents average of 2 technical replicates from one passage with 9 images taken per well. Clones E1, F6, and F8 were transduced with control GFP lentivirus, and clones C9, E11, and F3 were transduced with Pten GFP lentivirus. Significance was tested by an ordinary one-way ANOVA, with Tukey’s multiple comparison test. **J)** HGS2 subclones were grown in low-attachment u-bottomed plates for 90 hr. Each symbol represents the average of technical triplicates per subclone, performed at a different passage. The largest brightfield object area (µm^2^) per image was quantified and shown over time. **K)** HGS2 parental cells, control lentivirus or Pten lentivirus transduced subclones were injected I.P. into mice, and omental tumours harvested. Each symbol represents an individual mouse. Omental tumour weights are shown when harvested on days 56 (white) or 59 (filled). Control lentivirus clones F6 (triangle) and F8 (circle), and Pten lentivirus clones C9 (triangle), E11 (circle) and F3 (square). Significance was tested by one-way ANOVA, with Šidák’s multiple comparisons test. In all experiments, ****;p<0.0001, ***;p<0.001, **;p<0.01, *;p<0.05, ns=not significant.

Peritoneal dissemination is a key feature of HGSC, and tumour cells must resist anoikis and then grow in low attachment conditions to facilitate this dissemination. Both *Trp53*^-/-^ and *Trp53*^-/-^;*Pten*^-/-^ ID8 clones formed spheroid-like clusters in low-attachment *in vitro* (Fig. S1B), but contraction rates were equal over time in both genotypes (Fig. 1F). *In vivo, Pten* loss conferred no immediate survival advantage following intra-peritoneal injection (Fig. 1G; Fig. S1C). However, 14 days following injection, there was a significant expansion of *Pten* null cells in peritoneal fluid, suggesting that *Trp53*^-/-^;*Pten*^-/-^ ID8 cells utilise the microenvironment to enhance their growth (Fig. 1G). Moreover, tumour burden in the omentum, the dominant site of metastasis in HGSC, was greater by day 14, and significantly greater by days 25-28 (Fig. 1H) in mice injected with *Trp53*^-/-^;*Pten*^-/-^ cells.

To ensure our findings were not ID8-specific, we utilised HGS2, a cell line generated from tumours arising in a *Trp53^fl/fl^*;*Brca2^fl/fl^*;*Pten^fl/fl^*;*Pax8^Cre^*transgenic mouse model^24^. We re-expressed Pten in HGS2 using a lentivirus (Fig. S1D,E), which did not impact doubling time in 2D high-attachment (Fig. 1I) or the ability to form spheroids in low-attachment (Fig. 1J) but produced smaller omental tumours *in vivo* compared to control virus-infected cells (Fig. 1K). Together, these data suggest strongly that the peritoneal microenvironment supports accelerated growth of *Pten* null tumours.

We next hypothesised that macrophages support *Pten* null tumour seeding and growth. Two dominant macrophage populations exist in the peritoneal cavity and omentum in mice; F4/80^lo^MHCII^hi^ monocyte-derived macrophages, which are constantly replenished by blood Ly6C^hi^ monocytes, and embryonically-derived F4/80^hi^MHCII^lo^ resident macrophages, which are both self-maintained and replenished from the local F4/80^lo^MHCII^hi^ pool^25, 26^. Having confirmed the specificity of our gating strategy and our ability to distinguish macrophages from F4/80^+^SiglecF^+^ eosinophils (Fig. S2A-B), we first assessed how macrophage/monocyte populations altered during tumour growth. ID8 cell injection caused an influx of Ly6C^hi^ monocyte into peritoneal fluid within one day. This infiltration significantly increased in both peritoneal fluid and omentum by day 14 in *Trp53*^-/-^ ;*Pten*^-/-^-injected mice (Fig. 2A). An increase in F4/80^lo^MHCII^hi^ macrophages, likely to derive from this monocyte pool (Fig. 2B), was also increased 14 days after *Trp53*^-/-^;*Pten*^-/-^-injection. The resident F4/80^hi^MHCII^lo^ population only increased significantly at day 14 in *Trp53*^-/-^;*Pten*^-/-^-injected mice (Fig. 2C), which could result from local proliferation and/or *in situ* conversion of F4/80^lo^MHCII^hi^ macrophages. Interestingly on day 28, when both genotypes had substantial tumour burden, *Trp53*^-/-^ ;*Pten*^-/-^ omental tumours contained significantly more resident-like macrophages across multiple subclones (Fig. 2D). Furthermore, approximately 40% of resident-like macrophages expressed the long-term residency marker TIM4^+^, indicating that they are not newly recruited (Fig. 2E-F)^27^.

**Figure 2:**
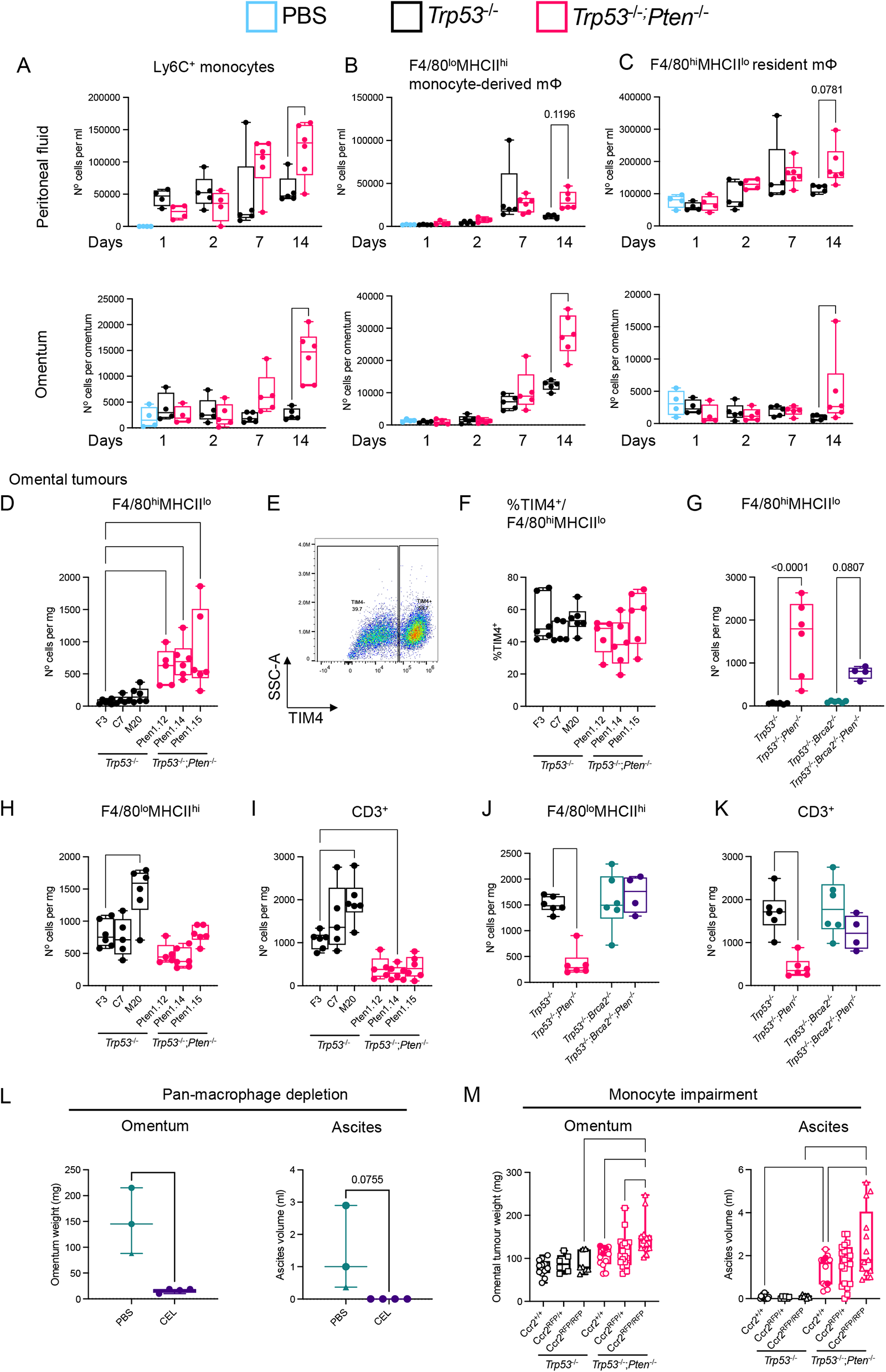
*Pten* null tumour cells enhance accumulation of resident-like macrophages within the omentum. **A-C)** Mice were injected with either ID8 *Trp53*^-/-^ (F3) or *Trp53*^-/-^;*Pten*^-/-^ (Pten1.14) cells. Peritoneal fluid and omenta were harvested 1, 2, 7 and 14 days later. Flow cytometry was performed for indicated cell populations. Counting beads were used to estimate absolute cell numbers, normalised to either the total lavage fluid (ml) or per omentum. Gating strategy for **A)** monocytes: Zombie Yellow^-^, CD45^+^, CD11b^+^, Ly6C^hi^. Macrophages: Zombie Yellow^-^, CD45^+^, CD11b^+^, Ly6C^-^, Ly6G^-^, SiglecF^-^, **B)** F4/80^lo^, MHCII^hi^ (monocyte-derived) or **C)** F4/80^hi^, MHCII^lo^ (resident-like). Every data point represents an individual mouse. Statistical significance was tested using a one-way ANOVA with Šidák’s multiple comparison test on selected samples. **D-F)** Mice were injected with individual ID8 *Trp53*^-/-^ (F3, C7 and M20) or *Trp53*^-/-^*;Pten*^-/-^ clones (Pten1.12, Pten1.14 and Pten1.15) on day 0 and omental tumours harvested at day 28 for flow cytometry. **D)** Resident macrophages were defined as Zombie Yellow^-^, CD45^+^, CD11b^+^, Ly6C^-^, Ly6G^-^, SiglecF^-^, F4/80^hi^, MHCII^lo^ and normalised to omental tumour weight (mg). **E)** Representative gating strategy used to define TIM4^+^ cells within the F4/80^hi^MHCII^lo^ population. **F)** Quantification of percentage TIM4^+^ cells out of total F4/80^hi^MHCII^lo^ macrophages. Statistical significance was tested using a one-way ANOVA with Šidák’s multiple comparison test on selected samples. **G)** Mice were injected with ID8 *Trp53*^-/-^ (F3), *Trp53*^-/-^*;Pten*^-/-^ (Pten1.14), *Trp53*^-/-^*;Brca2*^-/-^ (Brca2 2.14) or *Trp53*^-/-^*;Brca2*^-/-^;*Pten*^-/-^ (Brca2.14 Pten22) clones and the number of resident macrophages quantified by flow cytometry as in **D)**. **H)** Density of monocyte-derived, defined as Zombie Yellow^-^, CD45^+^, CD11b^+^, Ly6C^-^, Ly6G^-^, SiglecF^-^, F4/80^lo^, MHCII^hi^ cells in omental tumours of same mice as in **D)**. **I)** Density of T cells, defined as Zombie Yellow^-^, CD45^+^, CD3^+^ in omental tumours of same mice as in **D)**. **J)** Density of monocyte-derived, defined as Zombie Yellow^-^, CD45^+^, CD11b^+^, Ly6C^-^, Ly6G^-^, SiglecF^-^, F4/80^lo^, MHCII^hi^ cells in omental tumours of same mice as in **G)**. **K)** Density of T cells, defined as Zombie Yellow^-^, CD45^+^, CD3^+^ in omental tumours of same mice as in **G)**. **L)** Clodronate encapsulated liposomes (CEL) (n=6 mice) or PBS (n=3) were injected I.P. into mice on -14, -7 and -1 days prior to *Trp53*^-/-^*;Pten*^-/-^ (Pten1.12) tumour cell injection. CEL or PBS was then administered on days +7, +14, +21. Mice were harvested on day 26 (circles), apart from one PBS-treated mouse that reached endpoint at day 23 (triangle) and the omental tumour weight (mg) and ascites fluid volume (ml) was analysed. Statistical significance was tested using an unpaired t-test. **M)** *Ccr2*^+/+^, *Ccr2*^RFP/+^, *Ccr2*^RFP/RFP^ (clear symbols) or in-house wild-type (filled symbols) age-matched mice were injected with either *Trp53*^-/-^ (F3) or *Trp53*^-/-^;*Pten*^-/-^ (Pten1.14) ID8 cells on day 0 and culled on day 28. Omental tumour weight (mg) and ascites fluid volume (ml) were measured. Statistical significance was tested using a one-way ANOVA with Šidák’s multiple comparison test on selected samples (omental tumours) or with Tukey’s multiple comparison test (ascites). In all experiments, ****;p<0.0001, ***;p<0.001, **;p<0.01, *;p<0.05, ns=not significant.

To determine the impact of tumour cell *Pten* loss on resident macrophages further, we used *Trp53^-/-^ ;Brca2^-/-^* ID8 cells with or without *Pten* deletion. Loss of *Brca2* alone did not impact resident macrophage expansion (Fig. 2G). However, the additional deletion of *Pten* again significantly increased the density of resident macrophages (Fig. 2G), which was combined with a relative paucity of monocyte-derived macrophages and T cells (Fig. 2H-I). Deletion of *Brca2* in addition to *Pten* rescued the recruitment of monocyte-derived macrophages and T cells (Fig. 2J-K), suggesting that *Pten* deletion specifically alters resident macrophages rather than inducing global changes in the immune microenvironment (Fig. S3A-B).

To identify how *Pten* null tumour cells drive resident macrophage expansion, we screened chemokine and cytokine expression. This identified significantly increased *Ccl2* and *Ccl7* expression in some, but not all, *Pten* null cells (Fig. S3C-E), nor in cells additionally lacking *Brca2* (Fig. S3D). We also analysed the expression of retinoic acid producing enzymes *Raldh1*, *2* and *3* as peritoneal and omental resident macrophages are supported by retinoic acid, which drives their resident gene expression programme, including *Gata6*^28, 29^. However, *Raldh1* expression was not consistently altered by *Pten* deletion (Fig. S3F), whilst *Raldh2* and *3* expression was negligible in all cells. The canonical macrophage survival factor *Csf1* was also not significantly altered by *Pten* deletion (Fig. S3G). Furthermore, *Trp53*^-/-^;*Pten*^-/-^ cells did not demonstrate an enhanced ability to recruit bone marrow-derived macrophages (BMDM) (Fig. S3H), whilst deletion of *Ccr2* caused a reduction in BMDM recruitment to both genotypes (Fig. S3I). We also analysed *Il6* expression as *PTEN* deletion can drive IL6 production in prostate cancer^30^. IL6 gene expression was weak (C_T_ 34-37) with no statistical difference between clones (Fig. S3J) and IL-6 protein levels were undetectable by ELISA (data not shown). We also analysed *Vegfa*, the gene for vascular endothelial growth factor (VEGF) which is known to drive ascites formation^31^ but this was also unaltered by *Pten* loss (Fig. S3K). Taken together, these data suggest that one or more factors beyond Ccl2 and Ccl7 are produced *in vivo* that support resident macrophage recruitment and expansion, possibly acting indirectly through another cell type.

To determine if resident macrophages are drivers of *Pten* null omental tumour growth, we first depleted all macrophages using intraperitoneal injection of clodronate-encapsulated liposomes (CEL) prior to tumour inoculation (Fig. S4A). This pan-macrophage depletion completely prevented *Trp53*^-/-^;*Pten*^-/-^ omental tumour formation and ascites production (Fig. 2L). Unfortunately, due to high mortality rates observed following CEL injection, as previously reported^32^, we were prevented from performing further studies using CEL in our institution. To dissect macrophage contribution to *Pten* null tumour growth further, we utilised mice that lack either one (*Ccr2^RFP/+^*) or two copies (*Ccr2^RFP/RFP^*) of *Ccr2* and consequently have markedly reduced bone marrow monocyte egress^33^. As expected, monocytes and monocyte-derived macrophages were significantly reduced in the peritoneal fluid and omentum in both *Ccr2^RFP/+^* and *Ccr2^RFP/RFP^*mice, with no alteration in the resident macrophage pool following ID8 cell injection (Fig. S4B-E). Strikingly, deletion of *Ccr2* significantly increased tumour burden and ascites volume in *Trp53*^-/-^;*Pten*^-/-^-injected mice (Fig. 2M), indicating that the monocyte-derived macrophage pool has a protective anti-tumoural role during *Pten* null tumour seeding and growth.

Macrophage phenotypes cannot be simplified to binary M1/M2 marker expression^34^ and multiple distinct subtypes exist in omental tumours^35, 36^. Thus, we performed single-cell RNA sequencing on flow-sorted macrophages from omental tumours using the SMART-Seq2 protocol^37^ (Fig. S5A). UMAP clustering revealed five distinct clusters (Fig. 3A-B). Cluster 0 expressed genes classically found in monocyte-derived macrophages, including MHCII-associated molecules (*H2-Eb1, H2-DMb2, H2-DMb1, H2-Ab1, Cd74, H2-Oa, H2-Aa)*, chemokine receptors *Ccr2*, *Cx3cr1* and costimulatory molecule *Cd86* (Table. S2). Cluster 0 also localised in the F4/80^lo^MHCII^hi^ region by flow cytometry (Fig. 3C). Clusters 1 and 3 both expressed genes defined in peritoneal macrophages. Cluster 1 expressed *Fcna* (Ficolin 1), *Fn1* (Fibronectin 1)^38^ and the retinoid X receptor *Rxra*^39^ (Table. S2) and was predominantly located in the F4/80^lo^MHCII^hi^ region by flow cytometry (Fig. 3C). Cluster 3 expressed many more canonical peritoneal resident genes, including *Ltbp1*, *Garnl3*, *Serpinb2*, *Alox15*, *Selp*, *F5, Timd4*, *Icam2* (CD102) and *Gata6* (Table. S2)^38^. Cluster 3 was localized in the F4/80^hi^MHCII^lo^ region by flow cytometry (Fig. 3C), which, taken together with the expression of TIM4 (*Timd4*), suggests that cluster 1 is a monocyte-derived precursor that transitions into cluster 3. Cluster 4 expressed genes normally found in epithelial or mesothelial cells (*Krt18*, *Krt19*, *Msln, Wt1*), which suggests they may be phagocytic macrophages (Table. S2).

**Figure 3:**
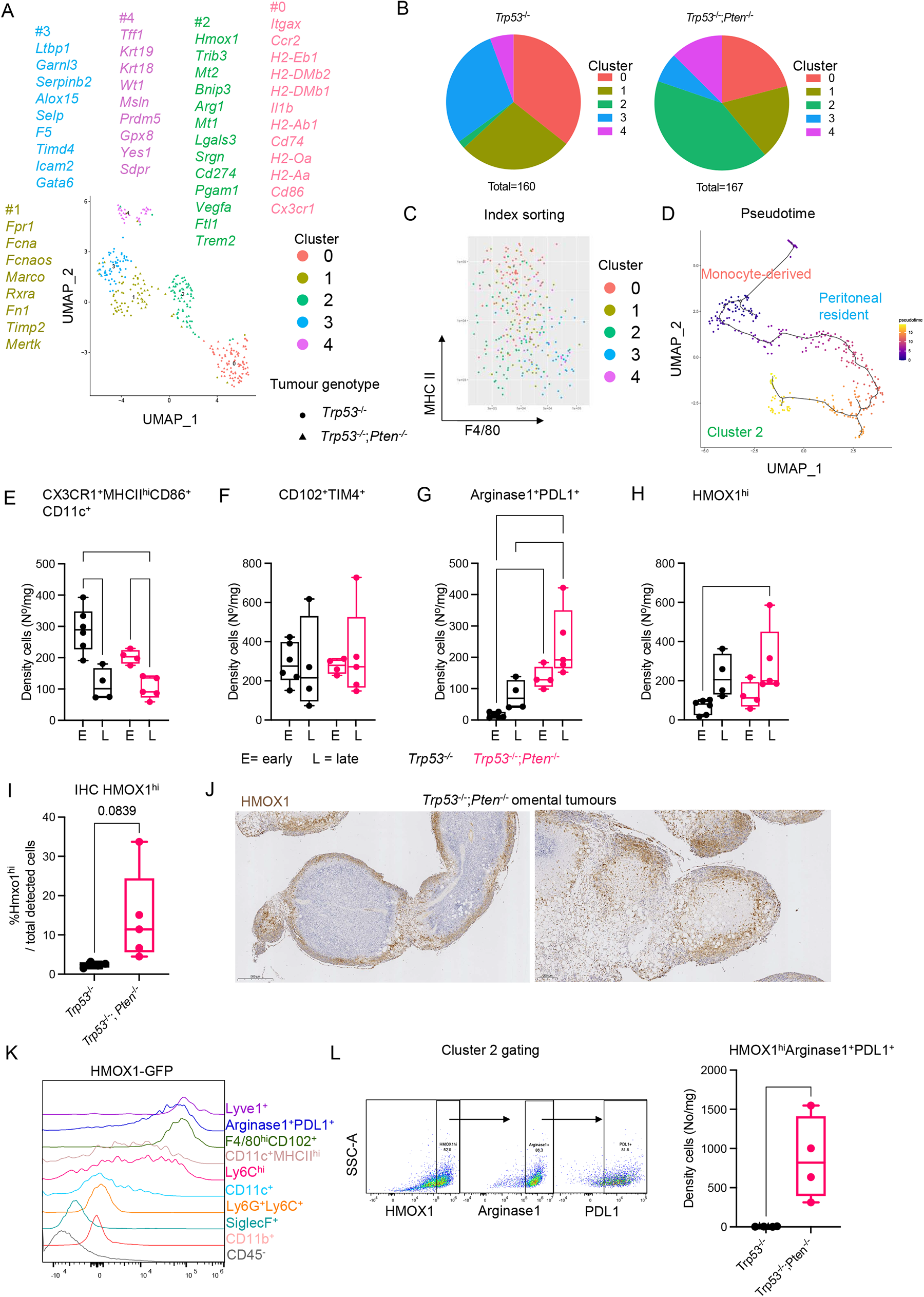
*Pten* null tumours drive accelerated formation of unique HMOX1^hi^ macrophage subpopulation. **A)** Mice were injected with ID8 *Trp53*^-/-^ (F3) or *Trp53*^-/-^;*Pten*^-/-^ (Pten1.14) ID8 cells on day 0 and omental tumours harvested at day 28, with n=4 mice per genotype. Macrophages were single-cell flow sorted based on DAPI^-^ (live), CD45^+^, CD11b^+^, Dump^-^ (CD3, CD19, Gr1), SiglecF^-^, F4/80^+^MHCII^+^, singlets and analysed by plate-based SMART-Seq2 single cell RNA sequencing. Following quality filtering, a UMAP projection of macrophages is shown, using Seurat pipeline. Selected significantly differentially expressed genes (DEGs) (defined as adjusted p value <0.05 and average log2-fold change >0) are shown next to respective cluster. Total DEGs per cluster are Cluster 0; 1022 genes, Cluster 1; 59 genes, Cluster 2; 425 genes, Cluster 3; 141 genes and Cluster 4; 1625 genes. **B)** The percentage of macrophages identified in each cluster isolated from either ID8 *Trp53*^-/-^ and *Trp53*^-/-^;*Pten*^-/-^ omental tumours from **A)** is shown. **C)** The expression of F4/80 and MHCII per macrophages, as collected during index sorting is shown with cluster identity overlaid by colour. **D)** Data in **A)** were reanalysed using the Monocle 3 package and Pseudotime analysis applied (shown as heatmap), with the root node placed in cluster 0. **E)** Mice were injected with ID8 *Trp53*^-/-^ or *Trp53*^-/-^;*Pten*^-/-^ ID8 cells on day 0 and omental tumours harvested at early (day 28 *Trp53*^-/-^; day 21 *Trp53*^-/-^;*Pten*^-/-^, “E”) and late (day 47 *Trp53*^-/-^; day 28 *Trp53*^-/-^;*Pten*^-/-^, “L”) timepoints. The density of macrophages in omental tumours was calculated for F4/80^+^MHCII^+^, CX3CR1^+^MHCII^hi^CD86^+^CD11c^+^. Statistical significance was tested by one-way ANOVA and Tukey’s multiple comparison test. **F)** As in **E)** the density of macrophages in omental tumours was calculated for F4/80^+^MHCII^+^, LYVE1^-^CD102^+^TIM4^+^. Statistical significance was tested by One-way ANOVA and Tukey’s multiple comparison test. **G)** As in **E)** the density of macrophages in omental tumours was calculated for F4/80^+^MHCII^+^, LYVE1^-^CD102^-^TIM4^-^Arginase1^+^PDL1^+^. Statistical significance was tested by One-way ANOVA and Tukey’s multiple comparison test. **H)** As in **E)** The density of macrophages in omental tumours was calculated for F4/80^+^MHCII^+^, HMOX1^hi^. Statistical significance was tested by One-way ANOVA and Tukey’s multiple comparison test. **I)** ID8 *Trp53*^-/-^ (F3) or *Trp53*^-/-^;*Pten*^-/-^ (Pten1.14) omental tumours harvested at day 28 were stained for HMOX1 by immunohistochemistry. The number of HMOX1^hi^ cells was quantified using QuPath. Statistical significance was tested using an unpaired t-test. **J)** Representative immunohistochemistry images of HMOX1 (brown stain) from *Trp53*^-/-^;*Pten*^-/-^ tumours from **I)** are shown, scale bar is indicated in the image (500 µm left image and 200 µm right image). **K)** HMOX1^GFP^ mice (n=2) were injected with ID8 *Trp53*^-/-^;*Pten*^-/-^ (Pten1.14) cells on day 0 and omental tumours harvested at day 25. Representative histogram of HMOX1-GFP expression is shown per cell population. The CD45^-^ population will contain transgenic stromal cells as well as the GFP^-^ ID8 cells. **L)** Gating strategy used to define cluster 2; F4/80^+^MHCII^+^, LYVE1^-^, CD11c^-^, MHCII^lo^, CD102^-^, F4/80^lo^, HMOX1^hi^, Arginase1^+^, PDL1^+^ (left). Density of cluster 2 macrophages in day 28 ID8 *Trp53*^-/-^ (F3) or *Trp53*^-/-^;*Pten*^-/-^ (Pten1.14) tumours (right). Statistical significance was tested using an unpaired t-test. In all experiments, ****;p<0.0001, ***;p<0.001, **;p<0.01, *;p<0.05.

The most interesting cluster was Cluster 2, which was found almost exclusively in *Trp53*^-/-^;*Pten*^-/-^ tumours (Fig. 3A-B) and had high expression of Heme oxygenase 1 (*Hmox1)*, an enzyme that catalyses the breakdown of heme into carbon monoxide, iron (Fe^2+^) and biliverdin. Cluster 2 also expressed genes involved in lipid accumulation, such as *Trib3* (Tribbles pseudokinase-3) and *Lgals3* (Galectin 3), as well as genes that protect against heavy metal toxicity, such as metallothioneins *Mt1* and *Mt2* (Table. S2). Cluster 2 also expressed genes associated with immunosuppression, including *Cd274* (PD-L1), *Arg1* (Arginase 1) and *Vegfa* (Table. S2). Cluster 2 localised mainly in the F4/80^hi^MHCII^lo^ region by flow cytometry, suggesting that it may derive from resident peritoneal macrophages (Fig. 3C). We performed pseudotime analysis (using Monocle3^40^) to estimate the putative direction of differentiation. When re-clustered (Fig. S5B) with cluster 0 selected as starting node, pseudotime predicted that cluster 2 derived from Clusters 1 and 3 (Fig. 3D), indicating that it represents a subtype of peritoneal resident macrophage.

We confirmed the presence of these macrophage subpopulations by flow cytometry at early (proceeding ascites formation) and late (ascites present) time points. Cluster 0, defined as CX3CR1^+^MHCII^hi^CD11c^+^CD86^+^, was significantly enriched in early *Trp53*^-/-^ tumours (Fig. 3E and Fig. S5C), correlating with our earlier data (Fig. 2H, J), but significantly declined with disease progression in both genotypes (Fig. 3E). We excluded LYVE1^hi^ cells, as they have been shown to define a separate mesothelial lining macrophage population^36^, whilst Clusters 1 and 3 formed a single peritoneal resident macrophage cluster defined as LYVE1^-^CD102^+^TIM4^+^. CD102^+^TIM4^+^ macrophages were abundant in both genotypes at both time points (Fig. 3F and Fig. S5D), indicating a steady expansion of resident peritoneal macrophages in the omentum during tumour growth, as reported in other settings^28, 39, 41^. Flow cytometry confirmed that Cluster 2 macrophages (defined as either Arginase1^+^PDL1^+^ or HMOX1^hi^) were present early in *Trp53*^-/-^;*Pten*^-/-^ tumours and increased significantly in late tumours (Fig. 3G-H and Fig. S5C-D). We validated this increased density of HMOX1^+^ cells in ID8 omental tumour sections using immunohistochemistry (Fig. 3I), where they were observed surrounding adipocytes and in tumour borders (Fig. 3J).

To understand if HMOX1 was a potential therapeutic target in HGSC, we first confirmed its selectivity for Cluster 2. Using *Hmox1*^GFP^ transgenic mice^42^, we confirmed that only monocytes and macrophages express HMOX1 (Fig. 3K). Although the LYVE1^+^ mesothelial lining population, which represent a small proportion of total macrophages, had the highest expression, cluster 2 (Arginase1^+^PDL1^+^) highly expressed HMOX1, followed by CD102^+^ peritoneal macrophages. CD11c^+^MHCII^hi^ monocyte-derived macrophages and monocytes had weak expression and other populations had weak/no expression. When combined, our data show that Arginase1^+^PDL1^+^HMOX1^hi^ macrophages are significantly enriched in *Trp53*^-/-^;*Pten*^-/-^ tumours (Fig. 3L).

To test the hypothesis that peritoneal fluid resident macrophages were the prime source of HMOX1^hi^ macrophages, we first adoptively transferred CD45.1^+^ peritoneal fluid cells (which will include monocytes, monocyte-derived and resident macrophages) into CD45.2^+^ mice 24 hr or 13 days following ID8 cell injection. CD45.1^+^ cells were detected in omental tumours, proving that trafficking can occur between peritoneal fluid and tumour (Fig. 4A). Interestingly resident CD45.1^+^F4/80^hi^MHCII^lo^ cells were enriched in *Trp53*^-/-^;*Pten*^-/-^ tumours (Fig. 4B-C, left panel), and correspondingly depleted in ascites (Fig. 4B-C, middle panel). The majority of CD45.1^+^ macrophages were TIM4^+^, indicating long-term residency (Fig. 4B-C, right panel). To determine further if HMOX1^hi^ macrophages can derive from peritoneal fluid, we sorted peritoneal fluid F4/80^hi^CD102^+^ cells from healthy HMOX1^GFP^ mice and adoptively transferred them into HMOX1^wt^ littermates bearing *Trp53*^-/-^;*Pten*^-/-^ tumours (Fig. 4D). We detected HMOX1^GFP^ cells in *Trp53*^-/-^;*Pten*^-/-^ omental tumours (Fig. 4E), which phenotypically copied the host’s own population (CD11c^-^MHCII^lo^, CD102^+^Arginase1^+^) and almost exclusively came from long-term resident CD102^+^TIM4^+^ cells (Fig. 4F). However, a fraction of CD102^-^ cells in the host macrophage pool had both high Arginase1 and PDL1 expression (Fig. 4G-H), suggesting that some Cluster 2 macrophages may also derive from non-CD102^+^ peritoneal fluid cells.

**Figure 4:**
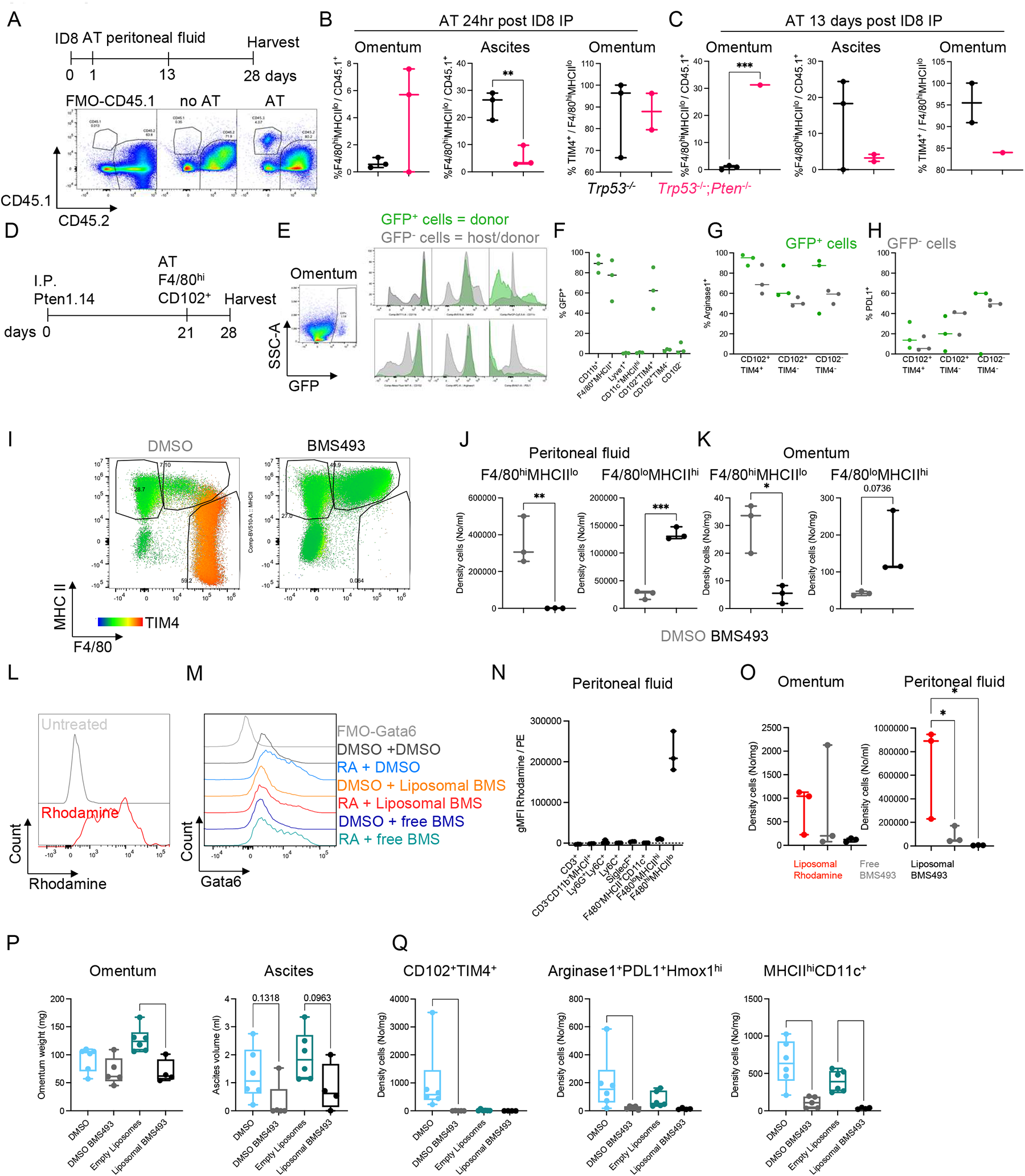
HMOX1^hi^ macrophages are partially derived from resident peritoneal fluid macrophages. **A)** CD45.2 mice were injected with ID8 *Trp53*^-/-^ (F3) or *Trp53*^-/-^;*Pten*^-/-^ (Pten1.14) cells on day 0 (n=6 per group). Mice then received an adoptive transfer (AT) of CD45.1 peritoneal fluid cells on either day 1 (n=3) or day 13 (n=3 for F3 and n=2 for Pten1.14) post ID8 I.P. Tumours and ascites were harvested at day 28. One mouse was excluded as there were insufficient cells and thus became a negative control. Representative flow cytometry gating strategy for live CD45.1 and CD45.2 cells in omental tumours. An FMO-CD45.1 control and no AT control are also shown. **B)** The omental tumours from **A)** were analysed by flow cytometry. The percentage resident F4/80^hi^MHCII^lo^ macrophages of all CD45.1 cells in omental tumour (left) and ascites (middle) are shown for mice that received CD45.1 AT 24hr post ID8 injection. The percentage TIM4^+^ out of F4/80^hi^MHCII^lo^CD45.1^+^ macrophages in omental tumour is also shown (right). One mouse had no detectable F4/80^hi^MHCII^lo^ macrophages, therefore the %TIM4^+^ value was not able to be analysed. Black values are from ID8 *Trp53*^-/-^ (F3)-injected mice and pink are from *Trp53*^-/-^;*Pten*^-/-^ (Pten1.14)-injected mice. Statistical significance was tested by unpaired t-test. **C)** The omental tumours from **A)** were analysed by flow cytometry. The percentage resident F4/80^hi^MHCII^lo^ macrophages of all CD45.1 cells in the omental tumour (left) and ascites (middle) are shown for mice that received CD45.1 AT 13 days post ID8 injection. The percentage TIM4^+^ out of F4/80^hi^MHCII^lo^CD45.1^+^ macrophages in omental tumour is also shown (right). Black values are from ID8 *Trp53*^-/-^ (F3)-injected mice and pink are from *Trp53*^-/-^;*Pten*^-/-^ (Pten1.14)-injected mice. Statistical significance was tested by unpaired t-test. **D)** F4/80^hi^ CD102^+^ peritoneal macrophages were FACS sorted from healthy *Hmox1*^GFP^ mice and adoptively transferred (AT) into *Hmox1*^wt^ littermates bearing ID8 *Trp53*^-/-^;*Pten*^-/-^ (Pten1.14) tumours on day 21. Omental tumours and ascites were harvested on day 28. **E)** Representative FACS plot of GFP^+^ cells in the CD45^+^ live gated cells in an omental tumour. The relative fluorescence GFP^+^ cells (green) compared to GFP^-^ cells (grey) is shown for markers CD11b, MHCII, CD11c, CD102, Arginase1, and PDL1. **F)** Macrophages were gated as previously, and the percentage of cells within each gate out of total detected GFP^+^ cells is shown. **G)** The percentage of each macrophage population gated within GFP^+^ (green) or GFP^-^ (grey) cells that is positive for Arginase1. **H)** The percentage of each macrophage population gated within GFP^+^ (green) or GFP^-^ (grey) cells that is positive for PDL1. **I)** Representative peritoneal fluid flow cytometry plot of F4/80 vs. MHCII (TIM4 is shown in pseudocolour) for healthy mice treated with BMS493 (20 mg/kg) on days 0, 2, 4 and culled on day 7. **J)** Density of F4/80^hi^ MHCII^lo^ (left) and F4/80^lo^MHCII^hi^ (right) macrophages in the peritoneal fluid of healthy mice following three doses of BMS493 (as in **I)**) compared to DMSO control. Statistical significance was tested using an unpaired t-test. **K)** Density of F4/80^hi^ MHCII^lo^ (left) and F4/80^lo^MHCII^hi^ (right) macrophages in the omentum of healthy mice following 3 doses of BMS493 (as in **I)**) compared to DMSO control. Statistical significance was tested using an unpaired t-test. **L)** BMDMs were incubated with medium or rhodamine-spiked liposomes for 3hr *in vitro* and their rhodamine fluorescence was analysed by flow cytometry. **M)** Peritoneal macrophages were seeded *in vitro* +/-1 µM retinoic acid. They were then treated with liposomal BMS493 or free BMS493 (or DMSO control) for 24hr and intranuclear Gata6 was analysed by flow cytometry. **N)** Healthy mice were treated with liposomal rhodamine I.P. on days 0, 3 and 5 and culled on day 7. The geometric median fluorescence intensity of rhodamine for each indicated immune cell population in the peritoneal fluid is shown. **O)** Healthy mice were treated as in **N)** with either liposomal rhodamine, liposomal BMS493 or BMS493 in DMSO. The density of F4/80^hi^ MHCII^lo^ macrophages in the healthy omentum (left) and peritoneal fluid (right) is shown. Statistical significance was tested for by one-way ANOVA and Tukey’s multiple comparisons test. **P)** Mice were treated with liposomal and free BMS493 twice-weekly starting on day -7, with *Trp53*^-/-^ (F3) or *Trp53*^-/-^;*Pten*^-/-^ (Pten1.14) ID8 cell injection on day 0. Twice weekly treatment was maintained until mice were culled on day 28. Omental tumour weight and ascites were quantified. Statistical significance was tested by ordinary one-way ANOVA with Šidák’s multiple comparisons test with selected comparisons. **Q)** The density of CD102^+^TM4^+^ (left), Arginase^+^PDL1^+^Hmox1^hi^ (middle) and MHCII^hi^CD11c^+^ (right) macrophages in omental tumours from **P)** are shown. Statistics by ordinary one-way ANOVA with Šidák’s multiple comparisons test with selected comparisons. In all experiments, ****;p<0.0001, ***;p<0.001, **;p<0.01, *;p<0.05, ns=not significant.

Peritoneal resident macrophages are dependent on retinoic acid^28, 39, 43^. We thus used BMS493, a pan-retinoic acid receptor (RAR) agonist that enhances the interaction of RAR with a nuclear co-repressor, preventing it from binding retinoic acid^44^. Treatment of healthy mice with BMS493 effectively depleted resident peritoneal macrophages and caused an expected recruitment of monocyte-derived macrophages, presumably to replace the niche of the resident population (Fig. 4I-K). To ensure RAR blockade was specific to macrophages, we encapsulated BMS493 in liposomes using the same formulation used for clodronate encapsulated liposomes^45^. Fluorescent, rhodamine-spiked liposomes were taken up rapidly by BMDMs *in vitro* (Fig. 4L), indicating that the liposomes could be phagocytosed by macrophages. Peritoneal macrophages pre-treated *in vitro* with retinoic acid express higher levels of the transcription factor Gata6, which was downregulated following liposomal BMS493 treatment (Fig. 4M). When injected into healthy mice, rhodamine-spiked liposomes were taken up specifically by F4/80^hi^MHCII^lo^ macrophages (Fig. 4N) and liposomal BMS493 effectively depleted F4/80^hi^MHCII^lo^ macrophages (Fig. 4O). We then treated mice with either free or liposomal encapsulated BMS493 before and after *Trp53*^-/-^;*Pten*^-/-^ ID8 cell injection. Treatment with liposomal BMS493 significantly reduced omental tumour burden and reduced ascites burden (Fig. 4P). As expected, both free and liposomal BMS493 effectively depleted CD102^+^TIM4^+^ macrophages in omental tumours (Fig. 4Q left) and Arginase1^+^PDL1^+^HMOX1^+^, expected to derive from them (Fig. 4Q middle). However, treatment with empty control liposomes also depleted CD102^+^TIM4^+^ macrophages (Fig. 4Q middle), which could be due to excessive lipid accumulation. Additionally, both free and liposomal BMS493 depleted MHCII^hi^CD11c^+^ monocyte-derived macrophages (Fig. 4Q right), which was not observed in our pilot experiments in healthy mice. This suggests that, during tumour growth, monocyte-derived macrophages also depend on retinoic acid or are indirectly impacted by the loss of peritoneal resident macrophages.

To overcome these limitations, we targeted HMOX1^hi^ macrophages directly using the HMOX1-inhibitor tin mesoporphyrin (SnMP)^46^. Treatment with SnMP did not impact the omental density of MHCII^hi^CD11c^+^ monocyte-derived, or CD102^+^F4/80^hi^ macrophages (Fig. 5A-B). However, SnMP treatment increased the density of Arginase1^+^PDL1^+^HMOX1^+^ macrophages (Fig. 5C). This is likely to be a compensatory mechanism in response to HMOX1 inhibition, as SnMP is known to stimulate HMOX1 upregulation whilst still blocking enzyme activity^47^. Interestingly, HMOX1 inhibition depleted LYVE1^+^ macrophages (Fig. 5D), which suggests they are susceptible to heme-induced toxicity, in line with their proximity to tumour vasculature^48^. Short term treatment with SnMP reduced ascites formation (Fig. 5E) although not omental tumour burden (Fig. 5F), whilst extended SnMP significantly increased survival of *Trp53*^-/-^;*Pten*^-/-^ tumour-bearing mice (Fig. 5G).

**Figure 5:**
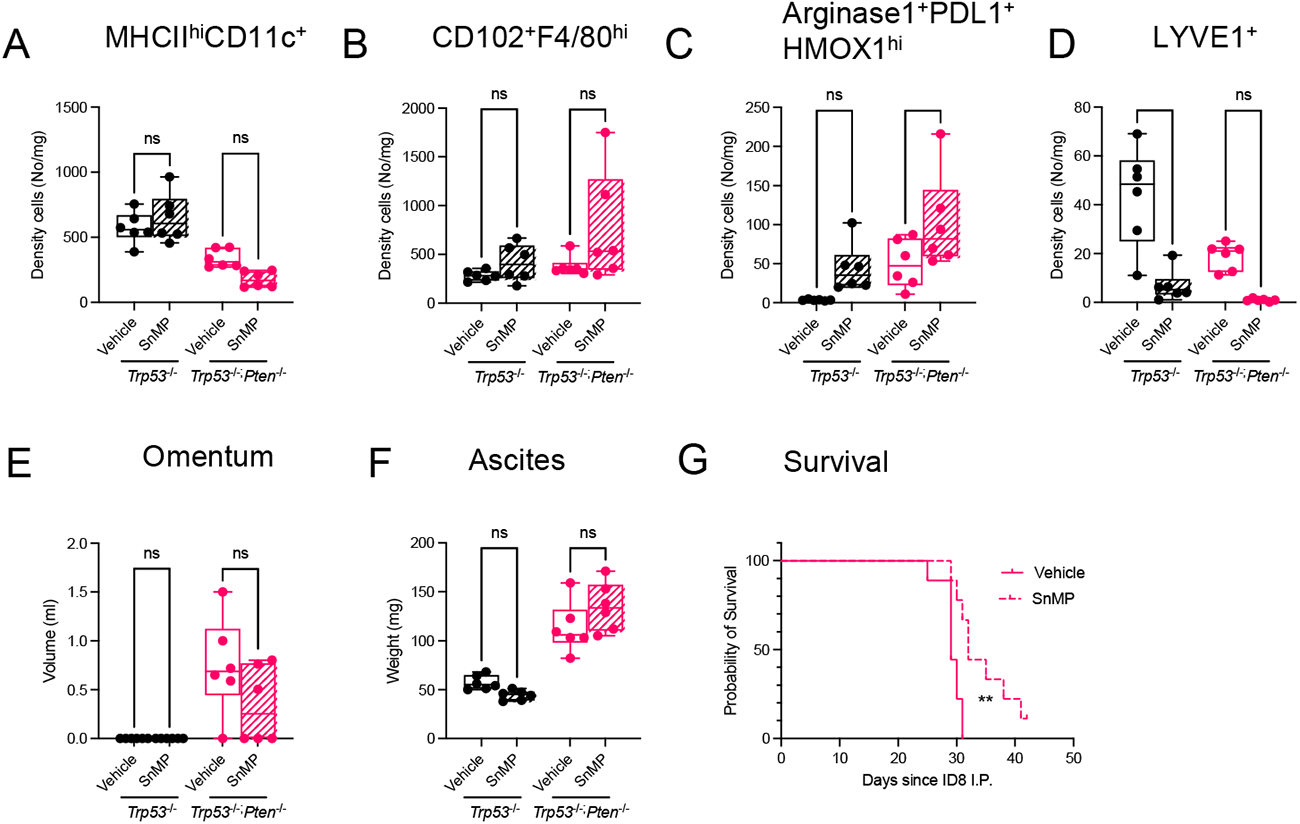
HMOX1 inhibition extends the survival in mice bearing *Pten* null ID8 tumours. **A)** Mice were injected with ID8 *Trp53*^-/-^ (F3) or *Trp53*^-/-^;*Pten*^-/-^ (Pten1.14) cells on day 0. From day 14, mice received 25 µmol/kg SnMP (n=6) or vehicle control (n=6) daily for 14 days. Omental tumours were harvested on day 28 and analysed by flow cytometry. The density of CD11c^+^MHCII^hi^ macrophages in the omental tumours is shown. Statistical significance was tested using one-way ANOVA and with Šidák’s multiple comparisons test with selected comparisons. **B)** The density of CD102^+^F4/80^hi^ macrophages from A) is shown per mg omental tumour. Statistical significance was tested using one-way ANOVA and Šidák’s multiple comparisons test with selected comparisons. **C)** The density of Arginase1^+^PDL1^+^HMOX1^+^ macrophages from A) is shown per mg omental tumour. Statistical significance was tested using one-way ANOVA and Šidák’s multiple comparisons test with selected comparisons. **D)** The density of LYVE1^+^ macrophages from A) is shown per mg omental tumour. Statistical significance was tested using one-way ANOVA and Tukey’s multiple comparison test. **E)** The omental tumour weight from A). Statistical significance was tested using one-way ANOVA and Šidák’s multiple comparisons test with selected comparisons. **F)** The ascites volume from A). Statistical significance was tested using one-way ANOVA and Šidák’s multiple comparisons test with selected comparisons. **G)** Mice were injected with ID8 *Trp53*^-/-^;*Pten*^-/-^ (Pten1.14) cells on day 0. From day 14, mice received 25 µmol/kg SnMP (n=9) or vehicle control (n=9) daily on a 5 days on/2 days off schedule until mice were harvested reached humane endpoint, which included advanced abdominal swelling. One mouse was censored in the SnMP group as it was killed before reaching endpoint at end of study (day 42); it had minimal disease present. Statistical significance was tested using a log-rank (Mantel-Cox) test. In all experiments, ****;p<0.0001, ***;p<0.001, **;p<0.01, *;p<0.05, ns=not significant.

To ensure our murine data were relevant to HGSC patients, we analysed a large single-cell RNA sequencing dataset, which contained data from 160 biopsies from 42 newly diagnosed, treatment naïve HGSC patients^49^. Tumour-associated macrophages (n=166,895) were annotated based on known marker genes including *PTPRC, CD14, FCER1G* and *CD68*. We first categorised human macrophages based on HMOX1 expression, defining those with a scaled HMOX1 expression >1 standard deviation above the mean as HMOX1^hi^ (Fig. 6A). We then compared differentially expressed genes (DEG) in HMOX1^hi^ macrophages with those for each mouse cluster. Cluster 2 showed the highest number of overlapping genes (n=39) with those in HMOX1^hi^ cells (Fig. 6B). Similarly, of the 87 DEG in HMOX1^hi^ macrophages, the highest proportion (44.8%, n=39/87) was shared with Cluster 2 (Fig. S6A). HMOX1^hi^ macrophages were enriched in key signature genes for resident macrophages (*LYVE1*), heme metabolism (*HMOX1*, *BLVRB*), cellular response to hypoxia (*HILPDA*), metallothioneins (*MT1E*, *MT1F*, *MT1G*, *MT1H*, *MT1M*), iron transporter, storage, and homeostasis (*SLC40A1*, *HAMP*, *FTH1*, *FTL*) and lipid metabolism and storage (*APOC1*, *PLIN2*, *LIPA*) (Fig. 6C, and Table S3). Conversely, HMOX1^lo^ macrophages were enriched in interferon gamma response genes (*CXCL9*, *CXCL10*), immune cell and T cell recruitment genes (*CCL5*, *CXCL9*, *CXCL10*, *CXCL11*, *IL1B*) and MHCII gene (*HLA-DQA1*) (Fig. 6C, and Table S3). MSigDB^50^ enrichment analysis of HMOX1^hi^ macrophage transcriptomes revealed an enrichment of hypoxia response, ion homeostasis, lipid metabolism and mTORC1 signalling pathways that were also found in mouse Cluster 2 (Fig. 6D, and Fig. S6B-C). Thus, human HGSC contains a cluster of macrophages that share common characteristics with mouse Cluster 2 and are characterised by high HMOX1 expression, tissue residency, oxidative stress response and low expression of pro-inflammatory cytokines/chemokines.

**Figure 6:**
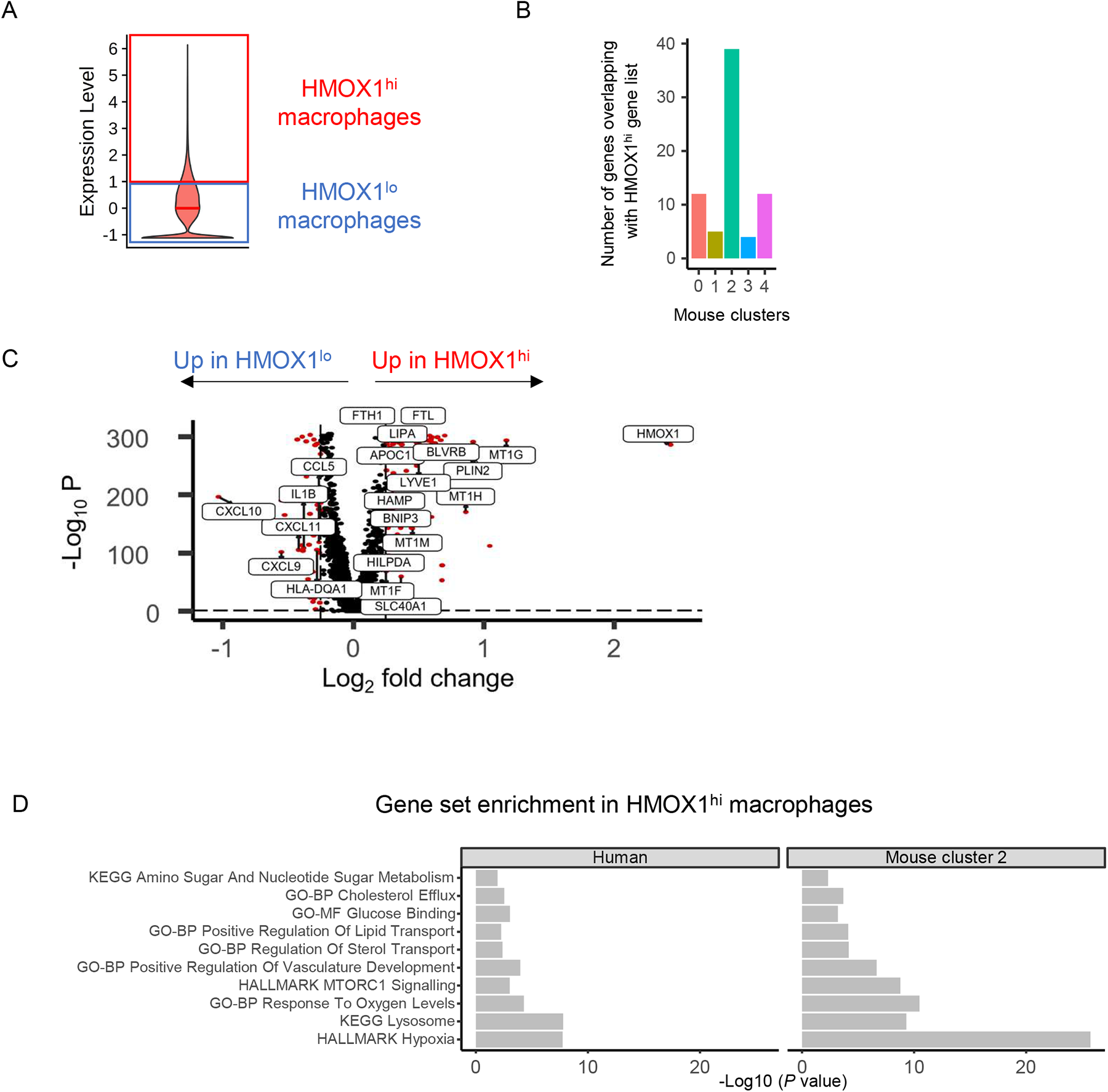
Mouse and human HMOX1^hi^ macrophages share common characteristics in HGSC. **A)** Scaled and centred HMOX1 expression on tumour associated macrophages. Macrophages with a scaled HMOX1 expression above 1 standard deviation from the mean were defined as HMOX1^hi^. Macrophages with a scaled HMOX1 expression below 1 standard deviation from the mean were defined as HMOX1^lo^. **B)** The overlap between DEG found in human HMOX1^hi^ macrophages and DEG found in each mouse macrophage cluster 0-4 is shown. **C)** DEG in HMOX1^hi^ macrophages (right side of the volcano plot) and HMOX1^lo^ macrophages (left side) defined in **A)** from human HGSC tumours is shown. **D)** Comparison of MSigDB pathway enrichment in human HMOX1^hi^ macrophages and mouse cluster 2 macrophages showing selected pathways of interest that were significantly enriched (Hallmark, Gene Ontology, KEGG).

In mice, Cluster 2 macrophages were found almost exclusively in *Pten* null tumours (Fig. 3A-B). In the single-cell RNA sequencing dataset, cancer cells from tumours enriched in HMOX1^hi^ macrophages (Fig. S7A) exhibited high mTOR signalling and high insulin-like growth factor signalling (Fig. 7A), which can activate PI3K/AKT signalling^51, 52^ (Fig. S7A). By contrast, tumours with a low proportion of HMOX1^hi^ macrophages were enriched in immune-related pathways (Fig. 7A). Immunohistochemistry on diagnostic HGSC samples from 172 patients in the BriTROC-1 study^53^ demonstrated a strong correlation between HMOX1 and CD68 (Fig. 7B-C), allowing us to use high HMOX1 expression as a surrogate for HMOX1^hi^ macrophages. The presence of HMOX1^hi^ macrophages positively correlated, albeit weakly, with positive *p*-AKT (S473) staining in tumour cells (Fig. 7D), a read-out for PI3K pathway activation^54, 55^ (Fig. 7E and Fig. S7B), and was also independently associated with reduced survival in BriTROC-1 patients, after adjustment for age and stage (HR = 1.80 [1.07-3.0]; Fig. 7F-G). The prognostic impact of high HMOX1 expression was confirmed in a separate validation cohort at the mRNA level (Fig. S7C).

**Figure 7:**
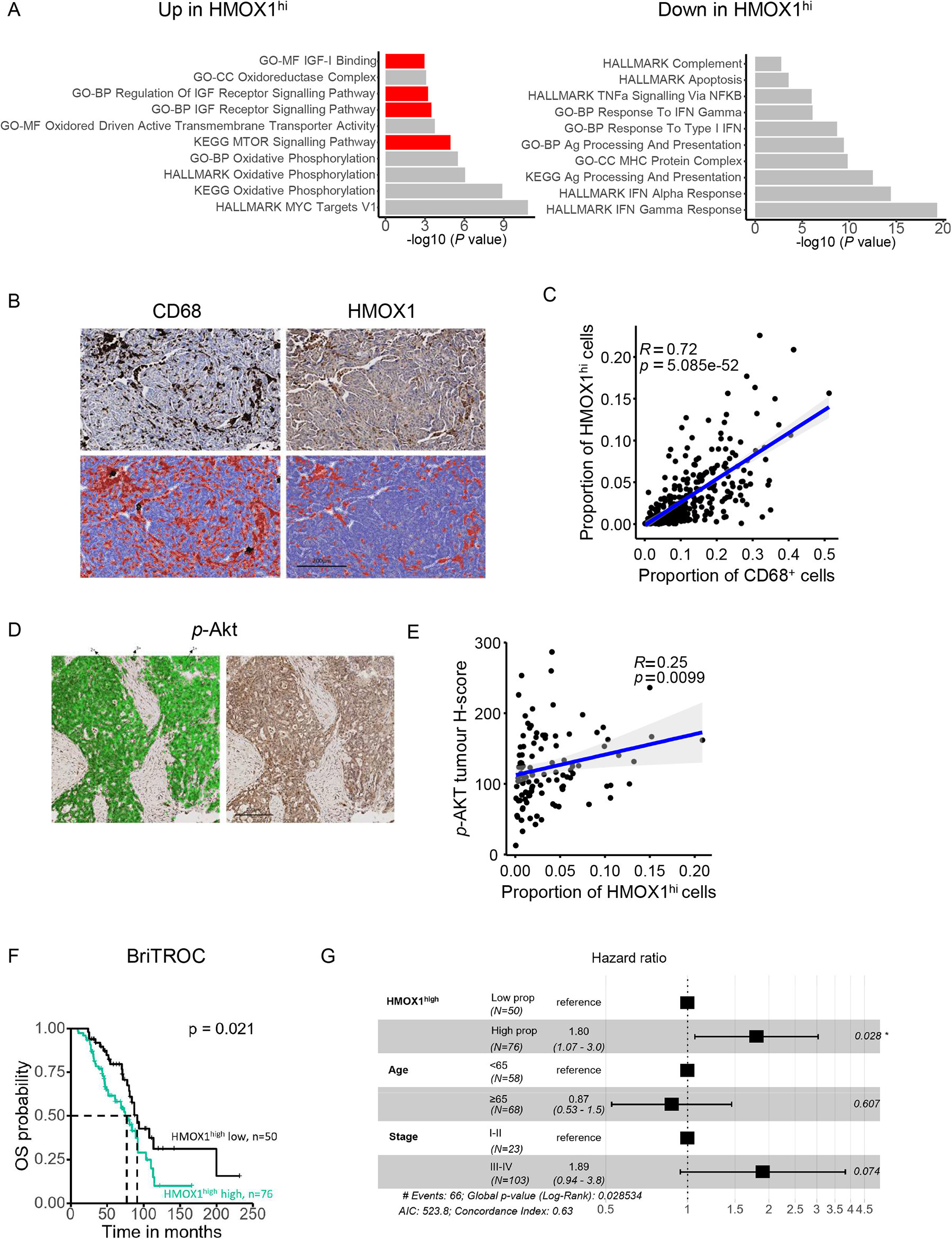
A high proportion of HMOX1^hi^ macrophages is associated with poor overall survival and PI3K signalling pathway activation. **A)** MSigDB enrichment analysis (Hallmark, Gene Ontology, KEGG) of HGSC tumours with high *vs* low proportion of HMOX1^hi^ macrophages showing selected pathways of interest that were significantly enriched (left) or downregulated (right). Pathways relating to PI3K-signalling are highlighted in red. **B)** CD68 (top left) and HMOX1 (top right) immunohistochemistry staining in the BriTROC-1 study TMA. QuPath positive cell detection is shown (in red) for CD68 (bottom left) and HMOX1 (bottom right). Scale bar represents 200 µm. **C)** Spearman correlation between the proportion of HMOX1^hi^ macrophages and the proportion of CD68^+^ macrophages found in BriTROC-1 TMA cores. **D)** *p*-AKT staining in the BriTROC-1 study with (left) and without (right) the QuPath tumour classifier showing weak (1+), moderate (2+) and strong (3+) staining. **E)** Spearman correlation between *p*-AKT tumour H-score and the average proportion of HMOX1^hi^ macrophages per patient in the BriTROC-1 study. **F)** Overall survival of patients in the BriTROC-1 study with high (n=76) and low (n=50) proportion of HMOX1^hi^, where the cut-off is based on the optimal threshold. Statistical comparison was performed using the logrank test. **G)** Multivariate regression forest plot of HMOX1^hi^ expression.

## Discussion

PTEN loss and other PI3K signalling alterations are frequent in HGSC but have proven challenging to target therapeutically. In this study, we have used mouse models and human HGSC samples to demonstrate that PI3K signalling pathway activation is associated with poor survival and the presence of HMOX1^hi^ macrophages. Importantly, we have shown that targeting this population with a specific HMOX1 inhibitor, SnMP, extends survival in mice. This suggests that targeting deleterious tumour-infiltrating macrophages has therapeutic potential.

Macrophages are the most abundant immune cell in HGSC^56^ but they have thus far eluded therapeutic targeting^57–59^. Multiple macrophage subtypes with diverse functions exist in HGSC^35, 36, 49^, and resident macrophages, derived from embryonic precursors^60^, dominate the pro-tumoural response^35, 36^. Conversely, we show here that monocyte-derived macrophages are protective against tumour growth in *Pten* null HGSC. This corroborates previous data in which anti-CSF1R treatment following carboplatin was shown to shorten survival via inhibition of the adaptive immune response^61^, and also the demonstration that stromal macrophage infiltration^62, 63^ and a high intratumoural HLA-DR:CD163 ratio correlates with improved survival^64^. Collectively, this indicates strongly that macrophage therapeutic approaches need to subtype specific.

Transcoelomic spread is the main mechanism by which HGSC disseminates around the peritoneal cavity and macrophages appear critical for this spread: gene expression in omental resident macrophages changes within hours of tumour cell injection in mice, whilst macrophages promote seeding of ID8 cells on the omentum, and macrophage depletion prior to tumour implantation prevents tumour seeding^65^. This early seeding is independent of T, B and NK cells, and occurs equally well in immunodeficient models^66^. Omental fat-associated lymphoid clusters (FALCs) are macrophage-rich, and resident embryonic-derived TIM4^+^CD163^+^ macrophages promote ID8 seeding and spread^35^. Additionally, omental-independent mesenteric-derived resident macrophages also support dissemination^36^.

*PTEN* loss and *PI3KCA* copy number alterations occur in HGSC carcinogenesis^67, 68^ and *Pten* deletion is essential for metastatic spread from the fallopian tube in transgenic murine models^69^ and also accelerates intraperitoneal tumour growth^22^. We show here that *Pten* deletion does not enhance proliferation or survival in low attachment conditions *per se*. However, *Pten* null cells specifically induce expansion of resident macrophages in the peritoneal fluid and omentum. Peritoneal resident macrophages support tumour spheroid formation and spread^70^ and targeting them via retinoic acid receptor deletion reduces tumour burden^39^. We show that *Pten* null tumours recruit peritoneal resident macrophages directly into the omentum, and that this is independent of blood monocyte recruitment, as *Ccr2*^RFP/RFP^ and *Ccr2*^+/+^ mice have equivalent omental resident macrophage numbers. This is important for considering targeting approaches, as recruitment does not occur directly from the blood. Furthermore, we show that *Pten* deletion accelerates formation of a unique resident macrophage population that expresses high levels of the heme-degrading enzyme, HMOX1.

HMOX1 expression is normally restricted to splenic and hepatic macrophages that remove senescent red blood cells, where its induction is cytoprotective against the oxidative stress induced by heme accumulation. However, aberrant expression in tumour associated macrophages drives immunosuppression^71, 72^, and HMOX1 inhibition by SnMP improves T cell infiltration and activity when administered with chemotherapy^71^. HMOX1 also induces expression of pro-inflammatory and angiogenic genes^73^, whilst HMOX1^hi^ macrophages can also directly drive metastasis, but not primary tumour growth, partly by aiding transendothelial migration and angiogenesis^74, 75^.

To target peritoneal resident macrophages, we first blocked retinoic acid signalling using liposomal BMS493, which significantly reduced tumour burden. However, targeting retinoic acid in the peritoneal cavity is likely to impact many cells in addition to macrophages^76^. In addition, BMS493 is lipophilic, and this unencapsulated BMS493 could not be purified from encapsulated when generating liposomes. Thus, liposomal BMS493 will also contain free drug that could generate off-target effects, and we were not confident liposomal BMS493 was specifically targeted resident macrophages.

However, crucially, we found that HMOX1 inhibition extended the survival of *Trp53*^-/-^;*Pten*^-/-^ ID8 tumour-bearing mice. SnMP is not directly cytotoxic to either tumour cells or macrophages^71^ and thus SnMP anti-tumoural activity is likely to be driven only via altered macrophage function. We found high HMOX1 expression in LYVE1^+^ macrophages^42^, previously shown to be mesenteric-membrane resident macrophages that can also promote ovarian cancer spread^36^. We did not detect this population in our single-cell RNAseq, most likely due to the small number of cells analysed. However, SnMP treatment did ablate LYVE1^+^ macrophages, whilst also driving an apparent increase in Cluster 2 macrophages. This reduction in LYVE1^+^ macrophages could result from their location in the perivascular niche and consequent susceptibility to heme-induced cytotoxicity^48^. This suggests that the more abundant Cluster 2 macrophages may upregulate HMOX1 in part by heme, but also by other microenvironmental factors, whilst future work will be required to determine whether the LYVE1^+^ population contributes in any way to the therapeutic effect here.

There are limitations in our study, not least that most of our findings derived from the ID8 murine model of HGSC, which is of ovarian surface epithelium origin^77^. However, ID8, and other OSE-derived models, such as STOSE^78^, can recapitulate the dominant features of HGSC, namely peritoneal dissemination and omental metastasis. However, it has been shown that different murine models represent the HGSC tumour microenvironment differently, as recently demonstrated^79^. For this reason, we attempted to replicate our findings using the fallopian-derived HGS2 line^24^. However, we could not stably restore wild-type *Pten* in HGS2 cells using CRISPR/Cas9 due to the *Brca2* deletion and consequent defective homology-directed repair^24, 69^. Nevertheless, omental tumour growth was reduced using HGS2 cells that had been transduced with a *Pten*-encoding lentivirus. This was not consistent across all clones tested, potentially due to promotor silencing, which is known to occur with lentiviruses^80^. Most importantly and reassuringly, two independent HGSC datasets validated our findings in mice: both the scRNAseq and IHC results reinforce the finding that the presence of HMOX1^hi^ macrophages is associated with poor outcome and activated PI3K signalling in HGSC.

Correlating murine and human data is extremely challenging^81^. Here we used Pten deletion to activate PI3K signalling in ID8 cells, whilst in HGSC, the pathway can be activated through multiple additional mechanisms, including *PIK3CA* and *AKT* mutation and amplification. We used *p-*AKT staining on IHC as a surrogate for pathway activation in patient samples, but it remains unclear whether every mechanism for activating PI3K signalling will generate HMOX1^hi^ macrophages. Similarly, the number of scRNAseq datasets available to interrogate tumour specific PI3K signalling remains small, and further data will be necessary to elucidate more nuanced biomarkers of pathway activity.

In summary, we have shown that HMOX1^hi^ macrophages, with common gene expression programmes including immunosuppression, hypoxia, cholesterol efflux, and lipid transport, can be identified in both murine and human HGSC. The function of HMOX1^hi^ macrophages in HGSC remains to be understood fully and the gene expression pathways may reflect both PI3K-driven microenvironment that induces HMOX1^hi^ cells and overall macrophage function. Nonetheless, our study highlights that HMOX1 inhibition may provide a relevant treatment strategy for HGSC.

## Figure Legends

**Supplementary Figure 1: Pten null cells are dependent on a tumour microenvironment for accelerated tumour growth.**

**A)** Representative biological replicate of *Trp53*^-/-^ (F3) or *Trp53*^-/-^;*Pten*^-/-^ (Pten1.14) ID8 cells seeded and grown in 4%, 0.4% or 0% FBS for 72hr, and imaged for 72 hr. Phase object count per well normalized to time 0 hr is shown.
**A)** Representative images of *Trp53*^-/-^ (F3) or *Trp53*^-/-^;*Pten*^-/-^ (Pten1.14) ID8 cells grown in low-attachment u bottomed plates, in triplicate, across 2 passages and imaged every 6hr for 168 hours (7 days). The largest brightfield object area (µm^2^) per image was quantified. A representative image of one replicate is show per cell line for each time point. Scale bar indicated on image, 800 µm.
**A)** Representative flow cytometry plots showing gating strategy to define ID8 cells in ID8-injected mice. ID8 cells were first gated on as CD45-, then by the viability dye Zombie Yellow and SSC-A high. Mice were injected with either PBS, ID8-F3 or ID8-Pten1.14 cells on day 0 and a peritoneal lavage was performed at days 1, 2, 7, and 14. Data from mouse 01, 05 and 09 from day 1 peritoneal lavages are shown.
**A)** FACs sorting strategy to purify HGS2 GFP^hi^ vs. GFP^neg^ as single-cells into a 96-well plate. HGS2 cells have been transduced with a control GFP or Pten GFP expressing lentivirus. FACS sorting strategy used to define GFP^hi^ cells, HGS2 cells were first gated on as GFP positive (P4) and the GFP^hi^ cells were further single cell sorted into 3x 96 well plates, 1 cell per well. GFP negative cells were also single cell sorted into 1x 96-well plate.
**A)** The single cell colonies generated in **D)** were visualised and grown into sublines, labelled by transduction and well of origin. Western blot was performed on HGS2 cells passaged 5 times since rethawing after single cell colonies were formed post transduction (P6 since conception). The membrane was cut above 70 kDa. The first 2 wells are ID8-F3 and ID8-TBK1 cells were used as positive controls for Pten expression. Pten protein is 54 kDa, GFP is 28 kDa, and β-Actin loading control is 42 kDa.
**A) L)** Representative images of lentivirus transduced HGS2 cells from **E)** were seeded in low attachment in technical triplicate and imaged every 6 hr for 7 days. Experiment was repeated across 3 passages. Scale bar indicated on image, 400 µm.

**Supplementary Figure 2: Defining resident and monocyte-derived macrophages in murine omental tumours.**

**A)** Digested omental tumours were analysed by flow cytometry and cell populations defined as follows: CD45^+^ (immune cells), Zombie^-^ (live), CD11b^+^ (myeloid), Ly6C^hi^ (monocytes), Ly6G^+^Ly6C^+^ (granulocytes), Ly6G-, Ly6C-, SiglecF^+^, F4/80^lo^ (Eosinophils), F4/80^hi^ MHCII^lo^ (resident-like macrophages), F4/80^lo^ MHCII^hi^ (monocyte-derived macrophages), F4/80^-^, MHCII^+^, CD11c^+^ (conventional dendritic cells 2, cDC2). Representative plots of macrophages in both *Trp53*^-/-^ and *Trp53*^-/-^;*Pten*^-/-^ omental tumours are also shown.
**B)** Cell populations (y axis indicated by colour) in **A)** were assessed for their fluorescence intensity (x axis) of markers CD64, CX3CR1, CSF1R, MerTK, CCR1, CCR2 and CCR3. The respective fluorescence minus one (FMO) control for each marker is also shown for the F4/80^hi^MHCII^lo^ and F4/80^lo^MHCII^hi^ populations.

**Supplementary Figure 3: *Pten* null resident macrophage accumulation cannot be completely explained by chemokine secretion.**

**A)** Mice were injected with individual *Trp53*^-/-^ ID8 CRISPR clones (F3, C7 and M20) or *Trp53*^-/-^*;Pten*^-/-^ clones (Pten1.12, Pten1.14 and Pten1.15) on day 0 and omental tumours harvested at day 28 for flow analysis. The density of monocytes, granulocytes, eosinophils and cDC2s in omental tumours are shown. Significance was tested for in the 6 clones using a One-Way ANOVA and Šidák’s multiple comparison test (to F3 control).
**B)** Mice were injected with individual ID8 CRISPR clones *Trp53*^-/-^ (F3), *Trp53*^-/-^*;Pten*^-/-^ (Pten1.14), *Trp53*^-/-^*;Brca2*^-/-^ (Brca2 2.14) or *Trp53*^-/-^*; Brca2*^-/-^ ;*Pten*^-/-^ (Brca2 2.14; Pten22) clones on day 0 and omental tumours harvested at day 28 for flow analysis. The density of monocytes, granulocytes, eosinophils and cDC2s in omental tumours are shown. Significance was tested for using a One-Way ANOVA and Šidák’s multiple comparison test (comparing *Trp53*^-/-^ to *Trp53*^-/-^*;Pten*^-/-^ and comparing *Trp53*^-/-^*;Brca2*^-/-^ to *Trp53*^-/-^*; Brca2*^-/-^ ;*Pten*^-/-^.
**C)** ID8 clones *Trp53*^-/-^ (F3 and C7) and *Trp53*^-/-^*;Pten*^-/-^ (Pten1.11 and Pten1.14) T*rp53*^-/-^*;Brca2*^-/-^ (Brca2 2.14 and Brca2 3.15) were analysed by a chemokine and cytokine gene array, in technical duplicates. Gene expression values were normalised to house-keeping genes as in methods. Volcano plot show the log2 fold change in gene expression relative to *Trp53*^-/-^.
**D)** Relative gene expression of *Ccl2* was measured by q-RT-PCR. *Trp53*^-/-^ (F3, C7 and M20), *Trp53*^-/-^*;Pten*^-/-^ clones (Pten1.12, Pten1.14 and Pten1.15), *Trp53*^-/-^;*Brca2*^-/-^ (Brca2 2.14) and *Trp53*^-/-^*;Pten*^-/-^;*Brca2*^-/-^ (Brca2 2.14; Pten22) ID8 clones were analysed. Each symbol represents the average of technical duplicates performed at different cell passages. Data analysed by 2∧(-ΔΔC_T_) method, using *Rpl34* as a housekeeping gene and F3 cells to normalise to as “1”. Raldh2 and Raldh3 isoforms were also analysed but had C_T_ values 35-40 and thus were deemed to be not expressed (data not shown). Significance was tested for in the 6 clones using a One-Way ANOVA and Šidák’s multiple comparison test (to F3 control). For the *Brca2* experiments, 2-3 technical replicates and 4 biological (different passage) replicates were performed. Every symbol represents the average of technical replicates performed at the same passage. One-Way ANOVA and Šidák’s multiple comparison test (*Trp53*^-/-^ vs. *Trp53*^-/-^*;Pten*^-/-^ and *Trp53*^-/-^;*Brca2*^-/-^ vs. *Trp53*^-/-^*;Pten*^-/-^;*Brca2*^-/-^).
**E)** Gene expression of *Ccl7 a*s in **D)**
**F)** Gene expression of *Raldh1 a*s in **D)**
**G)** Gene expression of *Csf1 a*s in **D)**
**H)** *Trp53*^-/-^ (M20) and *Trp53*^-/-^*;Pten*^-/-^ clones (Pten1.15) ID8 clones were seeded in a 24-well plate and the transwell migration of BMDMs assessed after 72hr coculture using crystal violet staining. Migration was normalised to a medium only control. Significance was tested using a one-way ANOVA and Tukey’s multiple comparison test.
**I)** *Trp53*^-/-^ (F3) and *Trp53*^-/-^*;Pten*^-/-^ clones (Pten1.14) ID8 clones were seeded in a 24-well plate and the transwell migration of BMDMs derived from either *Ccr2*^+/+^, *Ccr2*^RFP/+^, or *Ccr2*^RFP/RFP^ mice in a transwell assessed after 72hr coculture using crystal violet staining. Migration was normalised to a medium only control. Individual symbol shapes indicate BMDMs were derived from the same mouse. Significance was tested using a one-way ANOVA and Šidák’s multiple comparison test.
**J)** Gene expression of *Il6* was analysed as per **D)**, in *Trp53*^-/-^ (F3) and *Trp53*^-/-^*;Pten*^-/-^ (Pten1.14) ID8 clones. Each symbol represents 3 technical replicates performed across 3 passages. Data normalised to housekeeping gene *Rpl34*. Statistical significance was tested by an unpaired t-test.
**K)** Gene expression of *Vegfa* was analysed as per **D)**, in *Trp53*^-/-^ (F3) and *Trp53*^-/-^*;Pten*^-/-^ (Pten1.14) ID8 clones. Each symbol represents 3 technical replicates performed across 3 passages. Data normalised to housekeeping gene *Rpl34*. Statistical significance was tested by an unpaired t-test.

**Supplementary Figure 4: Depletion of macrophages alters *Pten* null ID8 tumour growth.**

**A)** Clodronate encapsulated liposomes (CEL) (n=6 mice) or PBS (n=3) were administered on days -14, -7 and -1 days prior to *Trp53*^-/-^*;Pten*^-/-^ (Pten1.12) tumour cell injection. CEL or PBS was then administered on days 7, 14, 21. Mice were culled on day 26 (circles), apart from one PBS-treated mouse that reached endpoint at day 23 (triangle) and the omental tumour and ascites was analysed by flow cytometry. The % of each population out of total CD45^+^ cells are shown. Gating strategy in as follows; Zombie-, CD45^+^, CD11b^+^, Ly6C^+^ (monocytes), Ly6G^+^ Ly6C^+^ (granulocytes), Ly6G^-^Ly6C^-^, SiglecF^+^ (eosinophils), F4/80^+^MHCII^+^(macrophages), or F4/80^-^MHCII^+^, CD11c^+^ (conventional dendritic cells 2). Significance was tested using Šidák’s multiple comparison test, comparing PBS to CEL for each site.
**B)** *Ccr2*^+/+^ (commercial wt mice), *Ccr2*^RFP/+^, *Ccr2*^RFP/RFP^ (clear symbols) or in house-bred wt (filled symbols) were age-matched and injected with either *Trp53*^-/-^ (F3) or *Trp53*^-/-^;*Pten*^-/-^ (Pten1.14) ID8 cells. Mice were culled on day 28 and the omental tumour was analysed by flow cytometry. The density per mg omental tumour is shown of **B)** monocytes **C)** F4/80^lo^MHCII^hi^ macrophages, **D)** F4/80^hi^MHCII^lo^ macrophages, **E)** cDC2, **F)** granulocytes, **G)** CD3^+^ T cells, **H)** CD4^+^ T cells, **I)** CD8^+^ T cell and **J**) CD19^+^ B cells. Data represents 4 independent experiments combined. Significance was tested using a one-way ANOVA and Šidák’s multiple comparison test.

**Supplementary Figure 5: Single-cell RNA sequencing analysis of omental macrophages**

**A)** Using the Seurat pipeline, the number of genes detected per cell (nfeature_RNA), number of reads per cell (nCount_RNA) and contribution of mitochondrial gene reads to total reads (percent.mt) are shown. Only nfeature_RNA that are detected in > 3 cells were considered, only cells with >200 nfeature_RNA and <10% mitochondrial gene expression (percent.mt) were accepted. F3T macrophages are from *Trp53*^-/-^ (F3) tumours and PTT macrophages are from *Trp53*^-/-^*;Pten*^-/-^ (Pten1.14) tumours. The number of the sample indicates the individual mouse samples used.
**B)** The Monocle 3 package was used to determine Pseudotime. Data was re-clustered by UMAP and the original cluster identity from Seurat pipeline shown by colour.
**C)** Flow cytometry gating strategy used to validate presence of single-cell RNAseq macrophage clusters in omental tumours. Total macrophages were first gated on as F4/80^+^MHCII^+^, then cluster 0 was defined as CX3CR1^+^MHCII^hi^, CD86^+^CD11c^+^. Cluster 2 was defined as CX3CR1^-^MHCII^lo^ and Arginase1^+^PDL1^+^. Fluorescence minus one (FMO) controls were used for some markers.
**D)** Flow cytometry gating strategy to validate presence of single-cell RNAseq macrophage clusters in omental tumours. Total macrophages were gated on as F4/80^+^MHCII^+^ as in **C)** then cluster 3 was defined as LYVE1^-^CD102^+^TIM4^+^. Cluster 2 was defined as LYVE1^-^ CD102^-^TIM4^-^HMOX1^hi^. Fluorescence minus one (FMO) controls were used for some markers.

**Supplementary Figure 6: Identification of HMOX1hi macrophages in human HGSC**

**A)** Pie chart showing overlaps in DEG between mouse macrophage clusters and HMOX1^hi^ macrophages set of genes (n=87).
**B)** MSigDB enrichment analysis of HMOX1^hi^ human macrophages showing top 20 pathways of interest that were significantly enriched (Hallmark, Gene Ontology, KEGG)
**C)** MSigDB enrichment analysis of mouse cluster 2 macrophages showing top 20 pathways of interest that were significantly enriched (Hallmark, Gene Ontology, KEGG)

**Supplementary Figure 7: Correlation of HMOX1^hi^ macrophages with HGSC disease**

**A)** Proportion of HMOX1^hi^ macrophages out of total macrophages per patient ranked in a descending order. Every I.D. on x axis represents an individual patient data. Selection of 10 patients with a high proportion of HMOX1^hi^ macrophages *vs* 10 patients with a low proportion of HMOX1^hi^ macrophages (black rectangles).
**B)** Spearman correlation between *p*-AKT tumour H-score and the proportion of HMOX1^hi^ macrophages per TMA core in the BriTROC-1 study.
**C)** Overall survival of HGSC patients (defined as serous ovarian cancers with a *TP53* mutation) with high and low expression of *HMOX1* in the KM plotter database, where the cut-off is based on the optimal threshold. Statistical comparison of survival curves was performed using the logrank test.

**Supplementary Table 1**: Table of antibodies used in flow cytometry and IHC.

**Supplementary Table 2**: Genes upregulated in each cluster from mouse clusters.

**Supplementary Table 3**: Complete list of up- and down-regulated DE genes in HMOX1^hi^ human macrophages.

## Methods

### Cell culture

Generation of ID8 cells with deletions in *Trp53, Pten* and *Brca2* has been described previously^21, 22^. ID8 cells were cultured in high-glucose 4.5 g/L DMEM (Life Technologies #21969035) supplemented with 4% heat-inactivated fetal bovine serum (Sigma and Life Technologies), 2mM glutamine (Life Technologies #25030024), ITS (Life Technologies #41400045) (10 µg/mL insulin, 5.5 µg/mL transferrin, and 6.7 ng/mL sodium selenite). HGS2 cells were purchased from Ximbio and were grown as previously described^24^ in DMEM:F12 Glutamax (Life Technologies #31331028), 4% FBS, murine epidermal growth factor (20 ng/ml) (Sigma #E4127) and Hydrocortisone (100 ng/ml) (Sigma #H0135), ITS. Early experiments were also performed 100 U/ml penicillin, 100 µg/ml streptomycin, 250 ng/ml Amphotericin B (Life Technologies 15240096) or 100 U/ml penicillin/100 µg/ml streptomycin. All cells were grown in 5% CO2, 37°C with humidity and used for a maximum of 10 passages. Cells were passaged using 0.1% Trypsin-EDTA (Gibco #15400054). Cells were regularly tested for mycoplasma using the Lonza MycoAlert^TM^ detection kit and were always negative.

### In vivo experiments

All *in vivo* work was performed at the Central Biological Services facility, Imperial College London in accordance with the U.K. Animals (Scientific Procedures) Act 1986 under Project Licences 70/7997, P2FEA2F22 and PA780D61A. Female C57BL6/J mice aged 6-7 weeks were purchased from Charles River, U.K. B6.129(Cg)-Ccr2tm2.1Ifc/J)^33^ (*Ccr2*^RFP/RFP^) mice were purchased from JAX (strain #017586). Both aged matched-control C57BL6/J mice and in-house bred WT mice were used as controls for *Ccr2*^RFP/RFP^ mice. Female 494C57BL/6L Y5.1 (CD45.1) mice aged 6-7 weeks were purchased from Charles River (strain #494) and used at 17 weeks. HO-1-Luciferace-eGFP-knock-in mouse (*Hmox1*^GFP^) mice were generated as previously^42^. All mice were acclimatised for at least 1 week prior to experiments. Mice were injected intraperitoneally with 1x10^6^ ID8 cells in 200 µl PBS or 10x10^6^ HGS2 cells in 200–300 µl PBS. Mice were monitored regularly and killed upon reaching moderate severity limit as permitted by the Project Licence limits, which included weight loss, reduced movement, hunching, jaundice and abdominal swelling.

### *In vivo* treatments

BMS493 (Bio-techne #3509/50) was prepared at 2.67 mg/ml in DMSO (5.2%) and sterile PBS and injected at 20 mg/kg I.P. In healthy mice BMS493 was injected on days 0, 2, 4 and culled on day 7. For liposomal BMS493 treatments see ***in vivo liposome treatments***. Sn(IV) Mesoporphyrin IX dichloride (SnMP) (Inochem #SNM321) was dissolved in 0.1M sterile NaOH and 0.5M NaHCO_3_ and injected I.P. at 25 µmol/kg. For fixed time point and flow cytometry analysis, mice were injected with ID8 cells on day 0 and SnMP was administered once daily (o.d.) from day 14 – 28, and mice were harvested on day 28. For the survival experiment, mice were injected with ID8 cells on day 0 and SnMP was administered o.d. from day 14 – 28, and mice were harvested upon when reaching humane endpoints. Mice with abdominal swelling but not yet at humane endpoint stopped receiving treatments (usually 1-3 days) before being culled to avoid bleeding.

### RNA extraction and cDNA synthesis

Cell medium was removed from 24-well plates and 350 µl RLT buffer added and frozen at -80°C. Plates were thawed on ice and 70% ethanol was added, gently mixed, and transferred into a RNeasy Micro Kit column (Qiagen #74104). RNA extraction was performed as per manufacturer’s instructions, including DNase step (Qiagen #79254). RNA was eluted in 30 µl nuclease free H_2_O and concentrated estimated using a Nanodrop. 2 µg of RNA was input into each 20 µl cDNA reaction using the High-Capacity cDNA Reverse Transcription Kit (Applied Biosystems #4368814) under cycling conditions 25°C 10 mins, 37°C 120 mins, 85°C 5 mins. The cDNA was diluted in 140 µl nuclease free H_2_O. qRT-PCR reaction was setup using 9 µl cDNA, 1 µl primer and 10 µl TaqMan Universal Master Mix II no UNG (Thermofisher #4440040). TaqMan primer probes were purchased from Thermofisher, *Rpl34* (Mm01321800_m1), *Ccl2* (Mm00441242_m1), *Ccl7* (Mm00443113_m1), *Csf1* (Mm00432686_m1), *Raldh1* (Mm00657317_m1), *Il6* (Mm00446190_m1), *Vegfa* (Mm00437306_m1). Samples were loaded in a 96-well plate (Applied Biosystems #4311971) and sealed with an Optical plate seal (Applied Biosystems #4346907) and analysed on a StepOnePlus (Applied Biosystems).

### SMART-Seq2 single-cell RNA sequencing

Briefly mice were injected with ID8 *Trp53*^-/-^ (F3) or *Trp53*^-/-^;*Pten*^-/-^ (Pten1.14) ID8 cells and omental tumours harvested at day 28, n=4 mice per genotype. 44 macrophages per tumour were flow sorted based on DAPI^-^ (live), CD45^+^, CD11b^+^, Dump^-^ (CD3, CD19, Gr1), SiglecF^-^, F4/80^+^MHCII^+^. SMART-

Seq2 single cell library preparation was performed by the Genomics Pipelines Group (Earlham Institute) and RNA sequenced on Illumina NovaSeq 6000 SP Lane (150bp paired end) with the aim for at least 1 million reads per cell. Further details on analysis provided in **Supplementary Methods**.

### CD45.1 adoptive transfer

CD45.2 mice were injected with ID8 F3 or Pten1.14 cells on day 0 (n=6 per group). Mice then received an adoptive transfer of peritoneal fluid cells either on day 1 (n=3 for both groups) or 13 days post I.P. (n=3 for F3 and n=2 for Pten1.14). A sterile peritoneal lavage (2 mM EDTA in PBS) was performed on two CD45.1 mice, samples combined and centrifuged at 330g 4 min 4’C. Cells were incubated with 5ml sterile RBC lysis buffer 5 min RT and washed in 15 ml PBS. 605,000 cells per I.P. were injected on day 1 and 400,000 cells on day 13. Omental tumours were harvested at day 28 and stained for flow cytometry.

### HMOX1^GFP^ macrophage adoptive transfer

A sterile peritoneal lavage (2 mM EDTA, 0.5% FBS in PBS) was performed on healthy *Hmox1*^GFP^ mice^42^and F4/80^hi^ CD102^+^ peritoneal macrophages were flow-sorted. 375,000 cells were adoptively transferred (AT) on day 21 into *Hmox1*^wt^ littermates previously injected with *Trp53*^-/-^;*Pten*^-/-^ (Pten1.14) on day 0. Omental tumours were harvested on day 28 for flow cytometry.

### Analysis of single-cell RNA sequencing (scRNA-seq) data

The data from a previous scRNA-seq study of 42 high grade serous ovarian cancer patients were analysed^49^ utilising the Seurat v4.3.0 R package^85^. HMOX1^hi^ macrophages were defined as macrophages^49^ that expressed *HMOX1* >1 standard deviation above the mean. The built-in FindMarkers function in the Seurat package was used to identify differentially expressed genes (DEG) and those adjusted p-values <0.05 were considered as differentially expressed. Adjusted p-values were calculated based on Bonferroni correction using all features in the dataset following Seurat manual [https://satijalab.org/seurat/v3.0/de_vignette.html]. Genes retrieved from Seurat analysis were displayed in a volcano plot using the enhancedVolcano package v1.14.0. MSigDB enrichment analysis of DEG between HMOX1^hi^ macrophages *vs* HMOX1^lo^ macrophages and between tumours with a high proportion of HMOX1^hi^ macrophages *vs* tumours with a low proportion of HMOX^hi^ macrophages was performed using msigdbr package v7.5.1 and clusterProfiler package v4.4.4 for Hallmark, Gene Ontology, and KEGG pathways.

### Prognostic value of HMOX1 in an independent validation cohort

The prognostic value of *HMOX1* mRNA expression was evaluated using the Kaplan–Meier Plotter (http://kmplot.com/analysis/)^90^. To analyse the overall survival (OS) of patients with HGSC (defined as ovarian cancer with serous histology and *TP53* mutation), we categorised the patients into two groups according to the best cut-off (high expression *vs.* low expression) of *HMOX1* (ID 203665_at) and assessed differences by using a Kaplan–Meier survival plot with hazard ratios, 95% confidence intervals, and log rank P values.

### Statistical analyses

A p value ≤0.05 was considered statistically significant. For immunohistochemistry, statistical analyses were performed in R (v4.2.1). HMOX1^hi^ expression was categorised using the optimal threshold from the maximally selected rank statistics (survminer package v0.4.9). Comparison of survival curves was performed using the logrank test. We performed COX multivariate regression of HMOX1^hi^ expression on clinical parameters such as age and FIGO stage and drew forest plot for visualization using the survival package v3.5.5. All other statistical analyses were performed using Prism v.9.4.1 (GraphPad).

## Acknowledgements

We thank the Imperial College London Central Biomedical Services for their expertise, advice, and assistance in performing all *in vivo* experiments. We also thank Dr Keira Turner for their help performing *in vivo* experiments. We also thank Dr William Jackson for their expertise and helpful discussion over results. We thank both the LMS/NIHR Imperial Biomedical Research Centre Flow Cytometry Facility and Department of Natural Sciences Flow Facility, Imperial College London for their help FACS-sorting and performing flow cytometry experiments. We thank Ignazio Puccio and Hiromi Kudo and the Section of Pathology, Department of Metabolism, Digestion and Reproduction, Imperial College London for the preparation of tissues for histology and IHC staining. We thank Dr Iain Macaulay and the Genomics Pipelines Group, Earlham Institute for their expertise and performing the SMART-Seq2 single cell sequencing. We thank Dr Nina Moderau, Department of Surgery and Cancer, Imperial College London for advice and access to reagents to perform lentivirus experiments. We also thank Dr Nazila Kamaly, Department of Chemistry, Imperial College London for advice in developing liposomes.

We are very grateful to all patients and their families that have provided samples that made this scientific research possible.

