## Supplementary Figures for "Heme oxygenase-1 expressing omental macrophages as a therapeutic target in ovarian high grade serous carcinoma"

**Supplementary Figure 1**

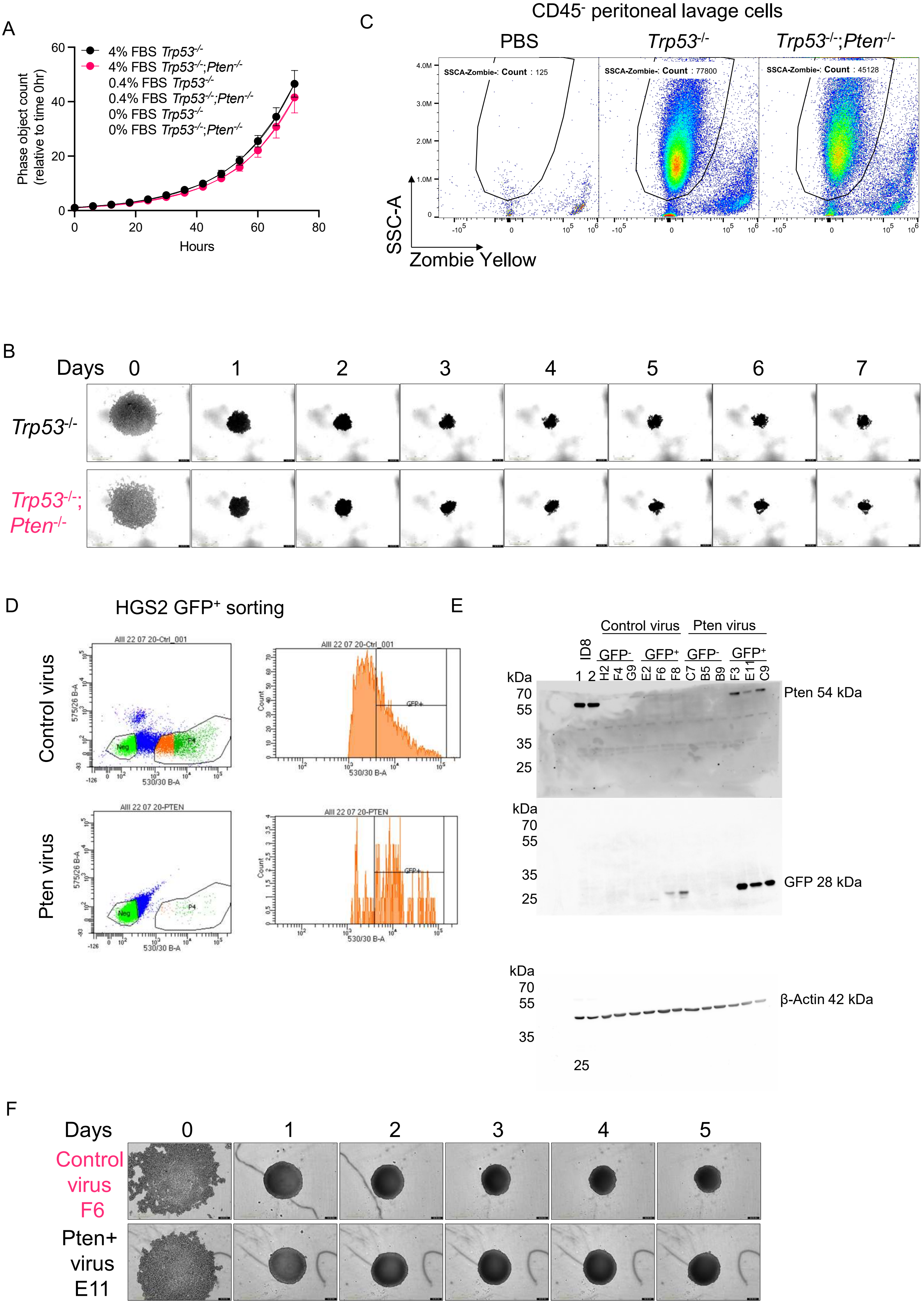

**Supplementary Figure 2**

A

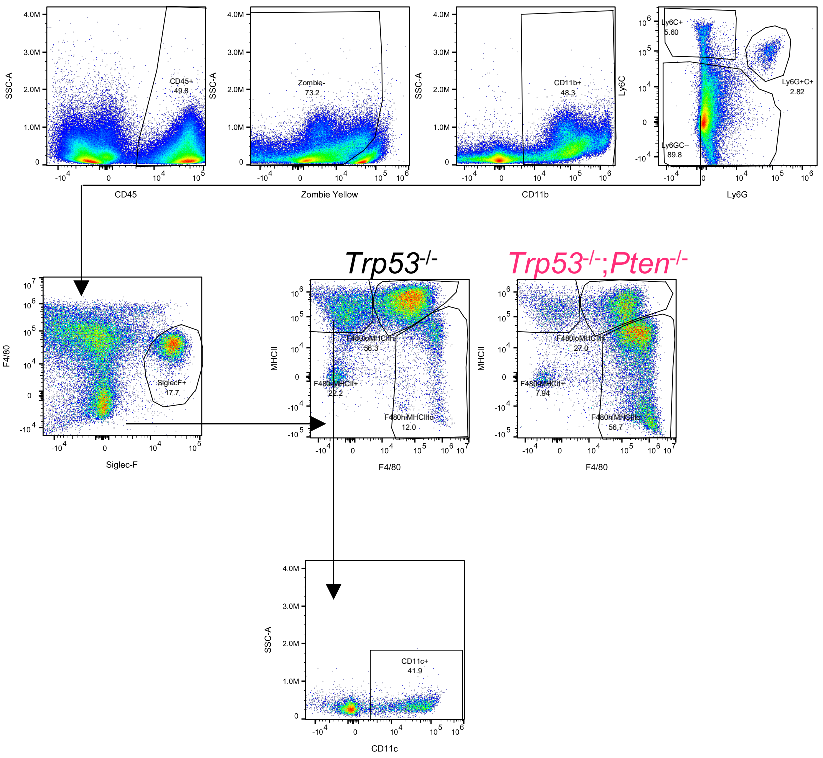


B

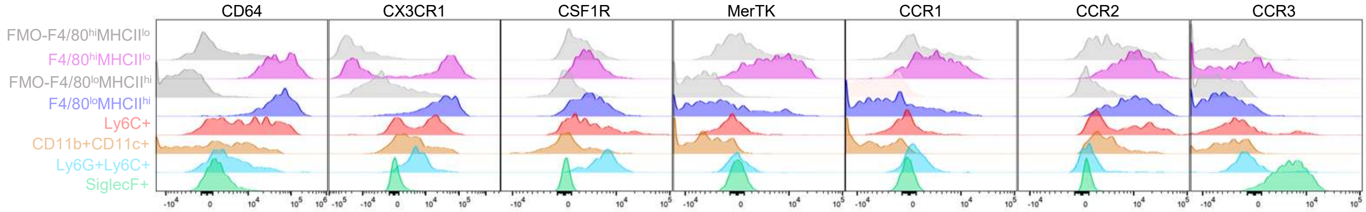

**Supplementary Figure 3**

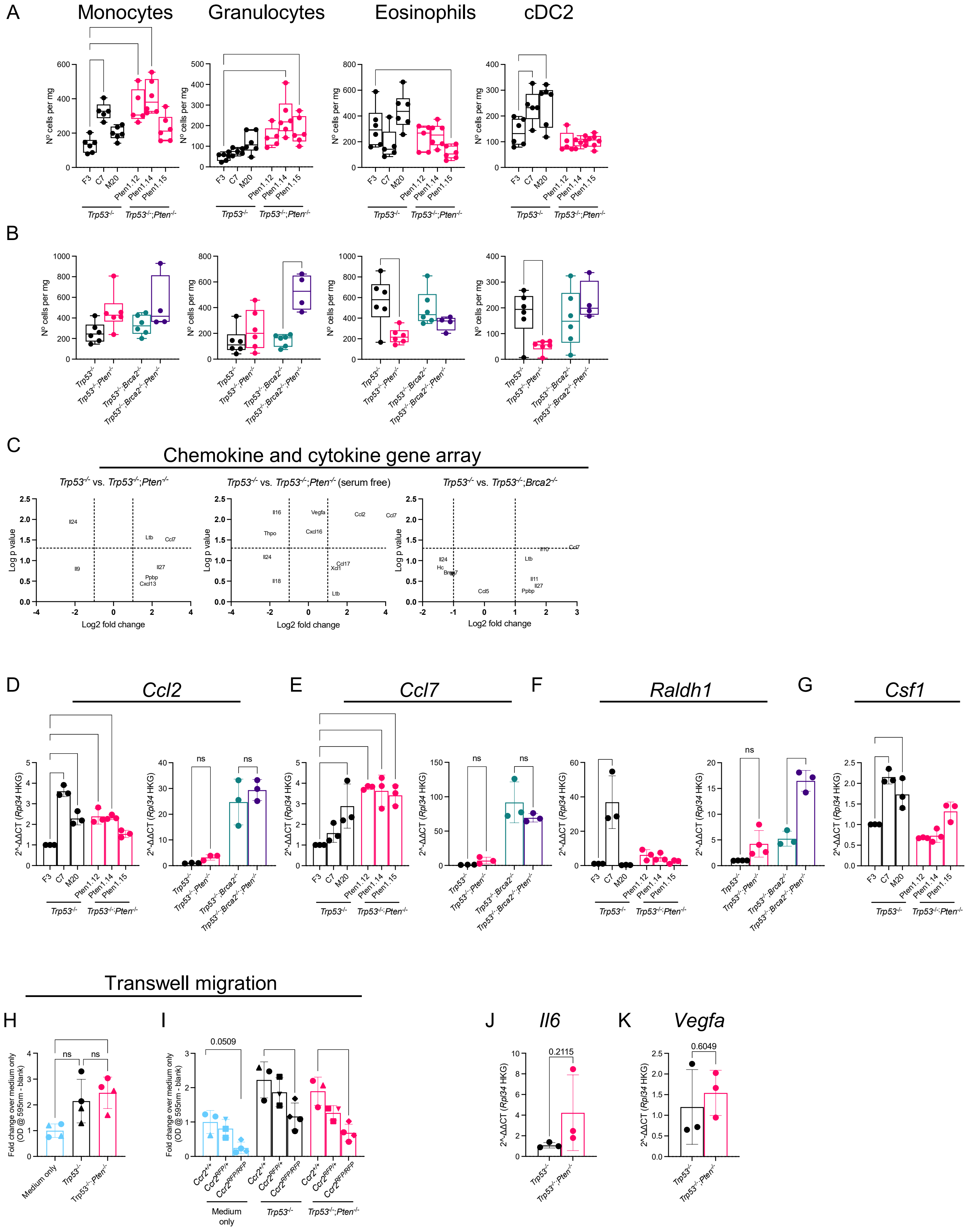

#### Supplementary Figure 4

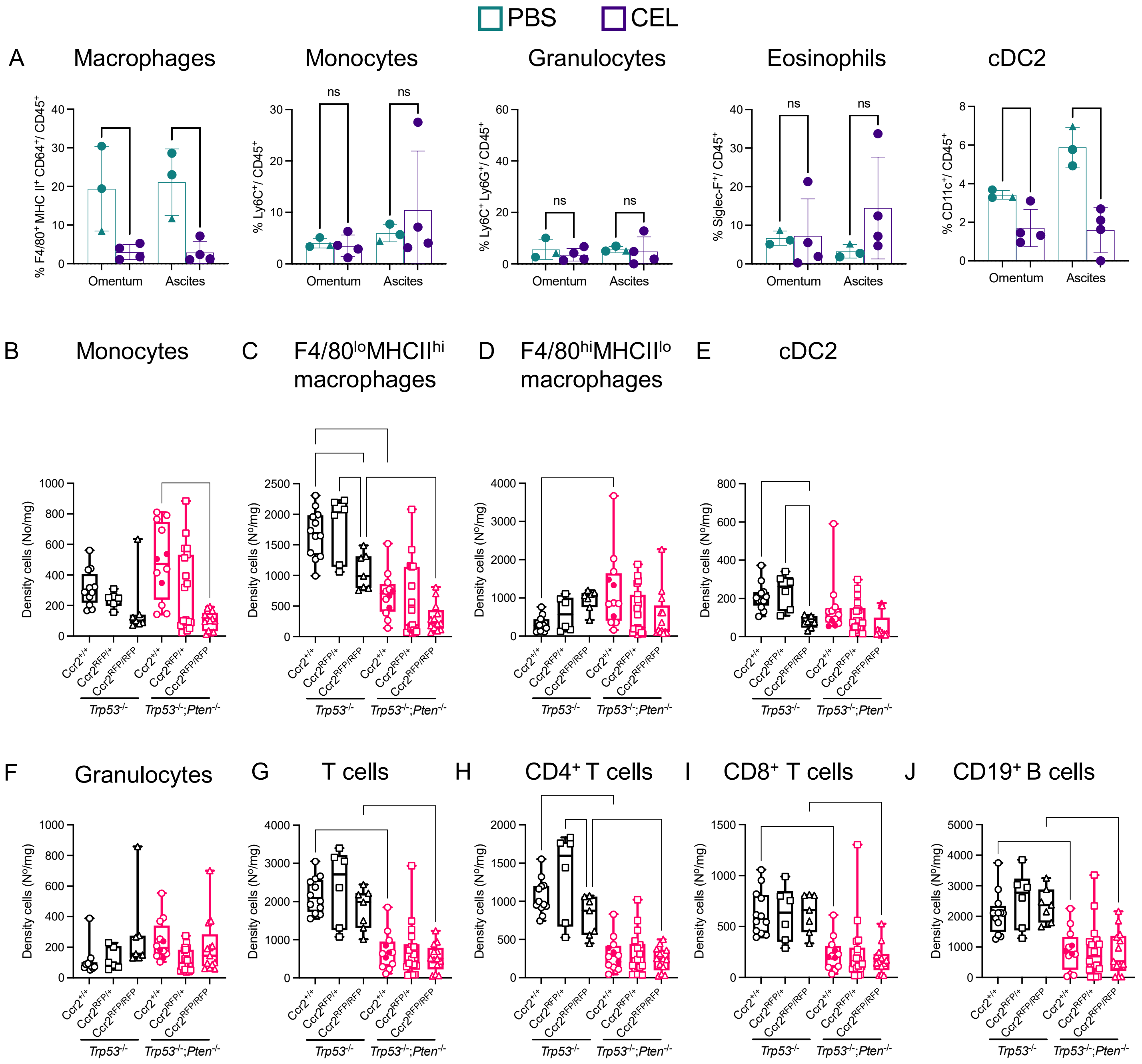

**Supplementary Figure 5**

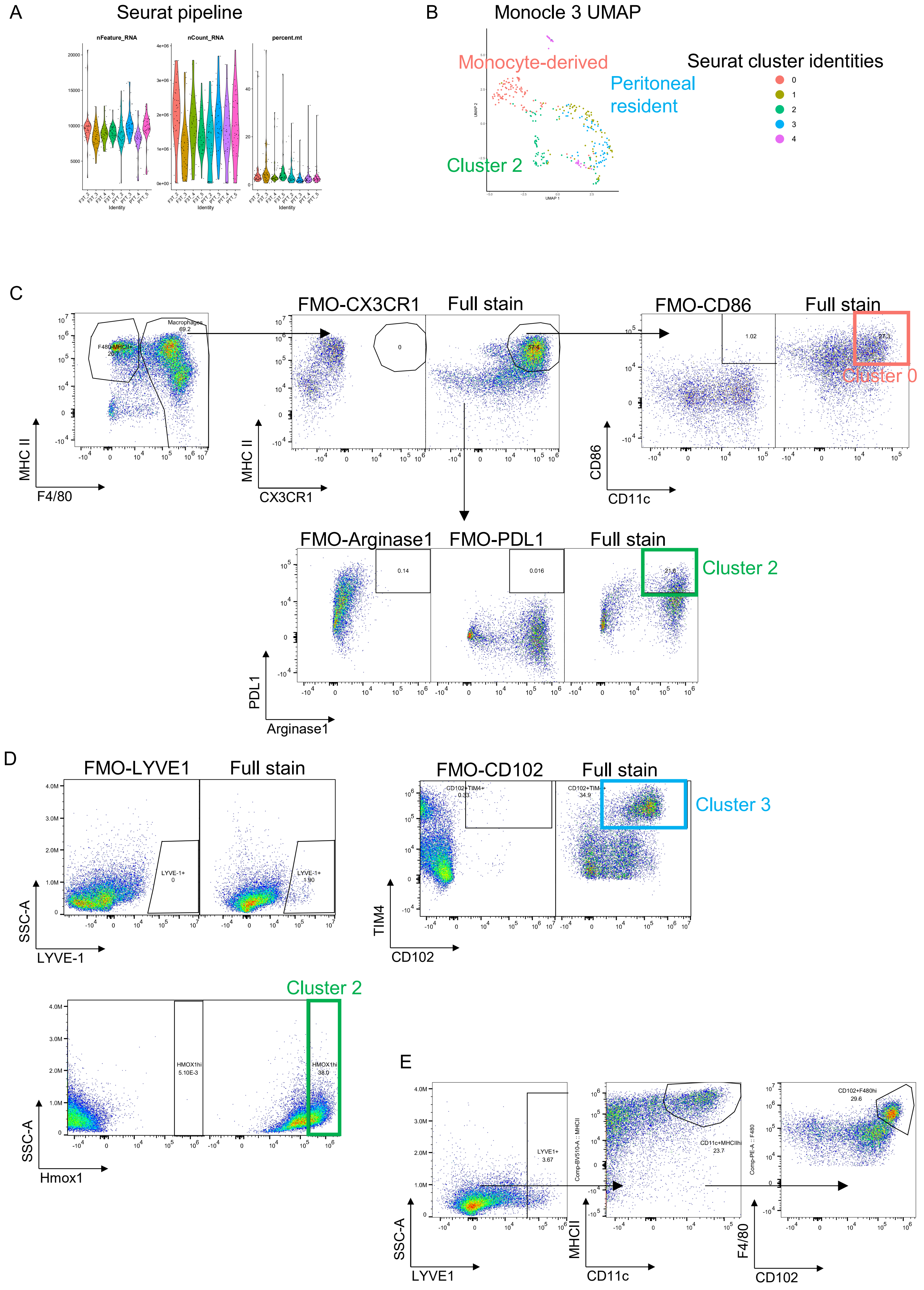

### Supplementary Figure 6

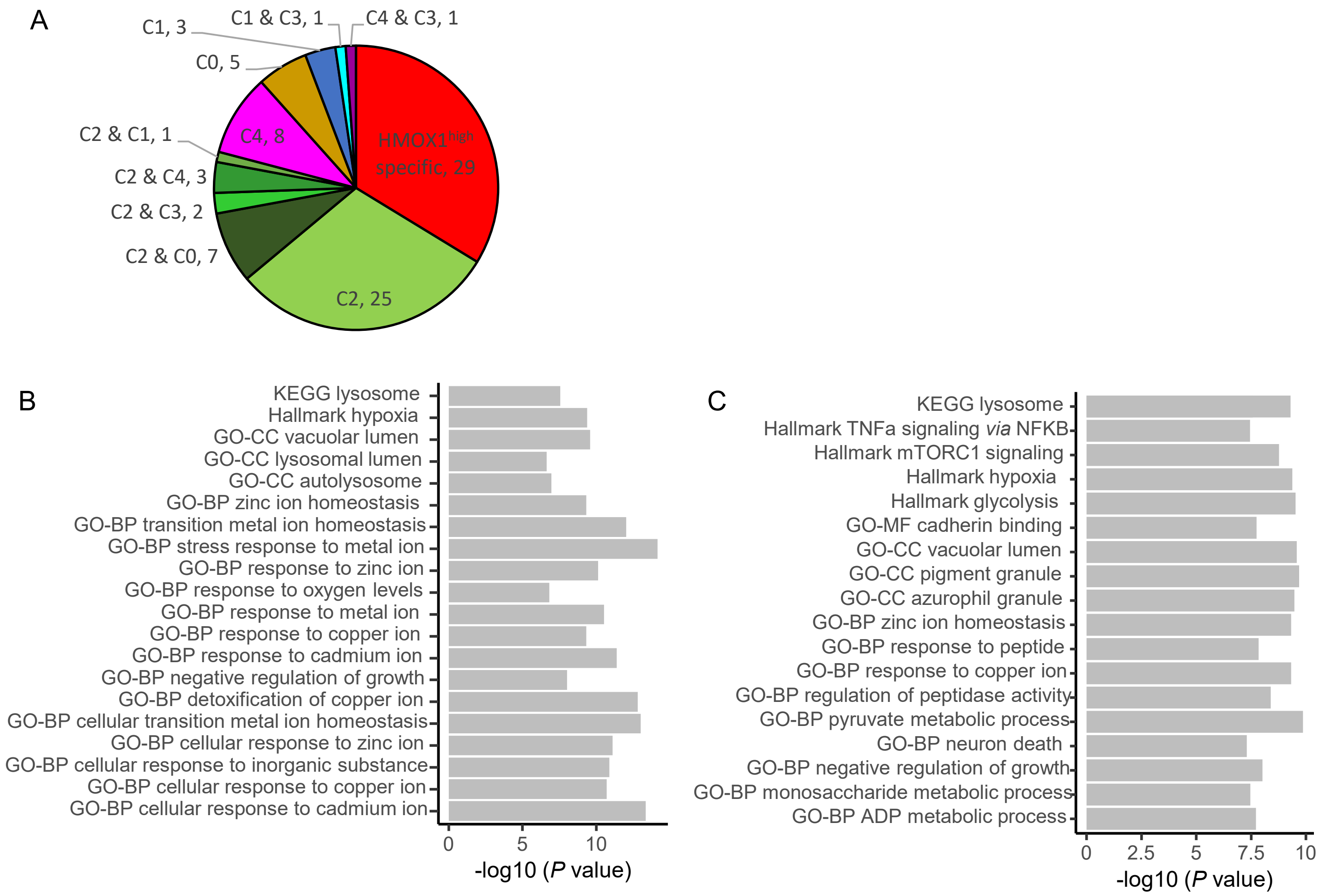

**Supplementary Figure 7**

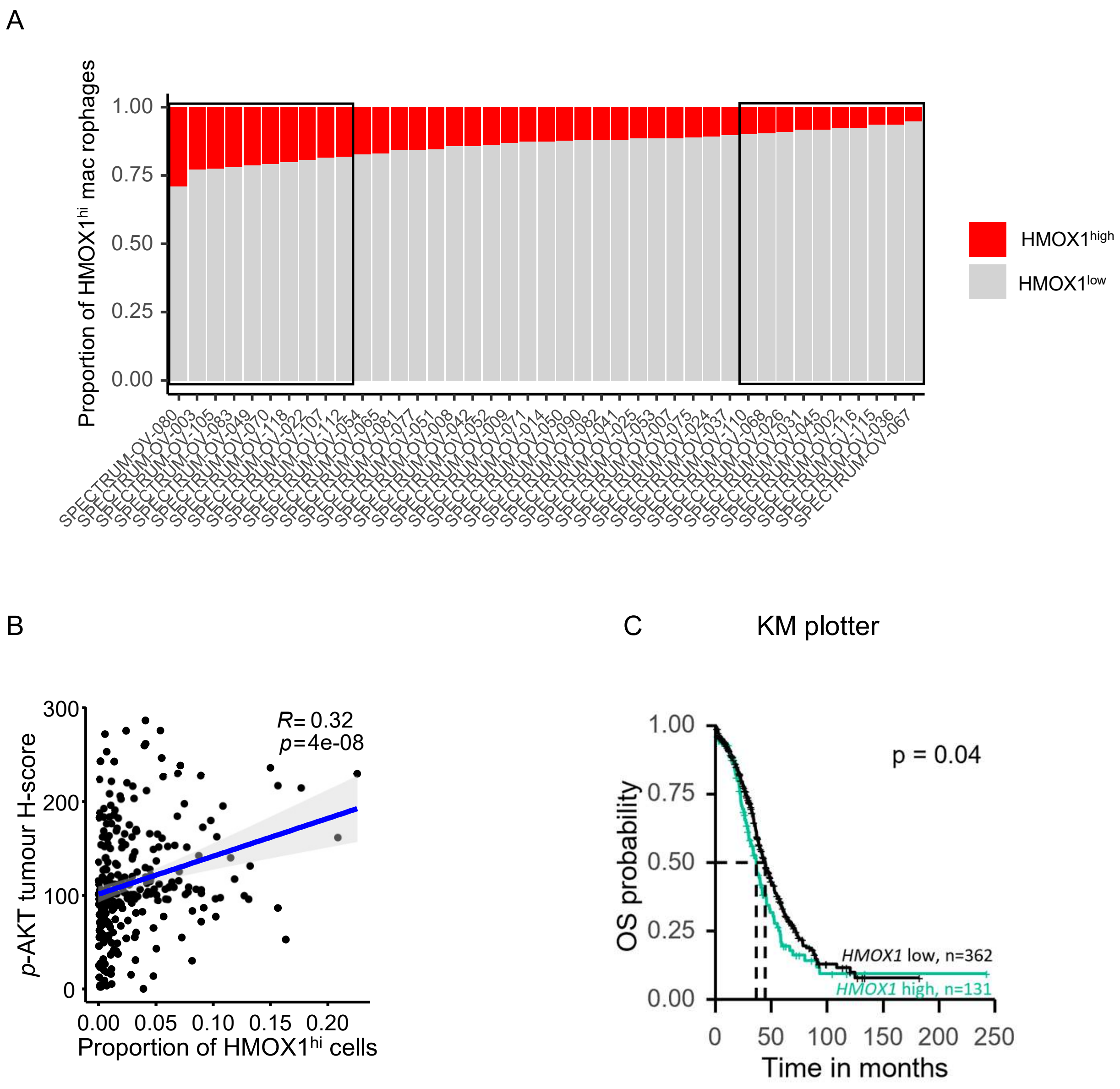
